# Membrane composition-dependent patterning of Rho and F-actin in an artificial cell cortex

**DOI:** 10.1101/2025.07.31.667950

**Authors:** Gregory J. Schwarz, Joanna R. Suber, Jennifer Landino

## Abstract

Cortical excitability, a phenomenon in which the cell cortex is dynamically patterned with waves of F-actin assembly, has been described in a variety of animal model systems, including embryos of mammals, flies, frogs and echinoderms, as well as a variety of cultured cells. While the cortical F-actin network is closely linked with the plasma membrane, it is not known if membrane composition or fluidity regulates dynamic cytokinetic patterning. Phospholipids partition within the plasma membrane during cytokinesis, and phosphoinositides play a key regulatory role in other excitable systems, suggesting a role for membrane-dependent regulation of cytokinetic patterning. Here we use an artificial reconstituted cell cortex comprised of *Xenopus* egg extract and supported lipid bilayers (SLBs) to show that membrane composition regulates self-organized cortical patterning. We find that manipulating levels of candidate lipids, including phosphatidylinositol 4,5-bisphosphate, phosphatidylethanolamine, sphingomyelin and cholesterol, results in both quantitative and qualitative changes in the dynamics of traveling waves and standing oscillatory patterns of active Rho and F-actin, as well as the kinetics of Rho activation and F-actin assembly on supported lipid bilayers. Our findings demonstrate that membrane composition directly regulates the assembly of cortical F-actin, as well as emergent active Rho and F-actin patterning.

**SIGNIFICANCE STATEMENT:**

- The cell cortex self-organizes dynamic patterns of active Rho and F-actin during cytokinesis, but it remains unknown whether and how the membrane composition impacts these dynamics.
- This study uses *in vitro* reconstitution of the cell cortex to directly manipulate membrane composition and finds that introducing different lipids induces changes in cortical dynamics of active Rho and F-actin.
- These findings reveal that membrane compositions regulates self-organized cortical dynamics, independently of changes to membrane fluidity. This work furthers our understanding of the mechanistic relationship between membrane composition, GTPase signaling, and cortical F-actin assembly.

## INTRODUCTION

The cell cortex is the outermost layer of the cell, and it is the primary determinant of cell shape in animals. The cortex comprises the fluid plasma membrane lipid bilayer and an adjacent cytoskeletal network made of filamentous actin (F-actin) and cross-linking proteins, which support the plasma membrane (Clark *et al*., 2013; Svitkina, 2020). Cortical F-actin drives changes in cell shape during essential functions like cell migration and cell division. Recent studies have shown that the cell cortex exhibits dynamic, periodic waves of F-actin assembly and disassembly in a variety of cell types and processes including cell adhesion (Case and Waterman, 2011; Barnhart *et al*., 2017), subcellular contraction (Graessl *et al*., 2017; Michaux *et al*., 2018; Yao *et al*., 2022), cell migration (Weiner *et al*., 2007; Gerhardt *et al*., 2014; Miao *et al*., 2017), mitosis (Xiao *et al*., 2017) and cytokinesis (Bement *et al*., 2015). Rho family GTPases drive this phenomenon, termed “cortical excitability”, and also exhibit dynamic patterning themselves (Bement *et al*., 2024). Rho GTPases act as molecular switches and, when activated, signal to downstream effectors that promote cortical F-actin assembly (Hodge and Ridley, 2016; Bement *et al*., 2024). Therefore, the subcellular localization and dynamics of Rho-family GTPases is critical for facilitating cortical remodeling and changes to cell shape.

Key to the function of Rho GTPases is their ability to directly interact with the plasma membrane. When active, Rho GTPases insert into the inner leaflet of the plasma membrane *via* C-terminal prenyl groups (Hodge and Ridley, 2016). They are spatially shuttled between the cytoplasm, where they are inactive and tightly bound to RhoGDI (Garcia-Mata *et al*., 2011), and the plasma membrane, where they are readily activated by membrane-localized guanine nucleotide exchange factors (GEFs) (Armstrong *et al*., 2025). The membrane localization of Rho GTPases - and thus Rho GTPase activity - is tightly regulated in cells, to ensure spatiotemporal control of downstream cytoskeletal dynamics. In cells, RhoA’s dwell time at the membrane is regulated by phosphoinositide-4,5-P_2_ (PIP_2_), through the scaffolding protein Anillin (Budnar *et al*., 2019). PIP_2_ is also thought to promote RhoA localization to the cell division site in a GEF-independent manner (Field *et al*., 2005; Wong *et al*., 2005; Yoshida *et al*., 2009; Liu *et al*., 2012). Notably, phospholipids within the plasma membrane are spatially redistributed during cell division, with PIP_2_, phosphatidylethanolamine (PE), sphingomyelin, and cholesterol enrichment at the cytokinetic furrow and midbody (Fernandez *et al*., 2004; Ng *et al*., 2005; Echard, 2008; Abe *et al*., 2012; Kunduri *et al*., 2022). *In vitro* studies have shown that phospholipids self-organize within a bilayer (Hansen *et al*., 2019) and in cells phospholipids can spatially oscillate within the plasma membrane (Gerhardt *et al*., 2014; Masters *et al*., 2016; Xiong *et al*., 2016). Additionally, phospholipid signaling has been shown to regulate cortical F-actin oscillations in non-dividing cells (Matsuoka and Ueda, 2018; Tong *et al*., 2023). Importantly, perturbations of lipid organization or metabolism during cytokinesis cause failed cell division and are associated with disease states including cancer (Kunduri *et al*., 2022). Therefore, defining the mechanistic relationship between membrane composition, phospholipid organization, and GTPase-mediated cortical patterning is essential to understanding cell functions like cytokinesis.

In developing *Xenopus laevis* and starfish embryos, cortical RhoA and F-actin waves are cell cycle-regulated (Swider *et al*., 2022) and are thought to support the contractile phase (C-phase, (Canman *et al*., 2000)) where the cortex is competent to form a cytokinetic furrow (Bement *et al*., 2015). These waves are supported by coupled positive and negative feedback signaling loops. Active RhoA recruits its own activator, Ect2 forming a positive feedback loop, and at the same time, drives F-actin assembly which recruits RGA3/4, a Rho inhibitor establishing negative feedback (Bement *et al*., 2015; Michaud *et al*., 2022). Remarkably, the core components of this excitability circuit are sufficient to induce cortical patterning in otherwise quiescent cells (Michaud *et al*., 2022). Our previous *in vitro* study, using *Xenopus laevis* egg extract, demonstrated that this excitable circuit drives self-organized patterning on a supported lipid bilayer (Landino *et al*., 2021). Notably, cortical patterns emerge from two populations of molecules: proteins and lipids. While previous work has focused on determining the mechanisms of protein signaling that support cortical excitability, here we investigate the less well-understood role of the membrane in regulating cortical pattern formation.

In this study, we leverage our reconstituted system, the artificial cortex, to investigate the role of membrane composition in regulating cortical patterning. The artificial cortex is comprised of *Xenopus laevis* egg extract and supported lipid bilayers (SLBs) and has been previously shown to reconstitute both traveling waves and standing oscillations of active Rho and F-actin (Landino *et al*., 2021). Traveling waves appear shortly after adding extract to SLB and are large-scale wave fronts that extinguish one another when they meet. Standing oscillations develop over time and have periodic association and dissociation with the SLB, but do not exhibit the spatial movement of traveling waves. These reconstituted patterns exhibit dynamics similar to those described *in vivo* (Bement *et al*., 2015; Landino *et al*., 2021), making this an ideal system to directly manipulate membrane composition and assay changes in cortical patterning in a manner that is not possible *in vivo*. Here, we found that active Rho and F-actin oscillations do not require cytoplasmic lipids or intracellular organelles. We also determine that key components of the physiological *X. laevis* oocyte plasma membrane (Hill *et al*., 2005), including PIP_2_, phosphatidylethanolamine (PE), sphingomyelin, and cholesterol, drive changes to cortex assembly and dynamics. Modifying the SLB composition alters the degree of F-actin assembly on the bilayer, the dynamics of traveling waves and standing oscillations, and the kinetics of active Rho and F-actin association on the bilayer. We therefore propose that membrane composition is a key regulator of self-organized cortical Rho and F-actin patterning during cytokinesis.

## RESULTS

### PE-containing SLBs alter traveling active Rho and F-actin waves

To investigate the role of membrane composition in regulating cortical patterning, we directly altered the composition of the SLB in the artificial cortex. Previous characterization of active Rho and F-actin dynamics in the artificial cortex used bilayers that were 60% PC, 30% PS, and 10% PI (Nguyen *et al*., 2014; Landino *et al*., 2021). We referred to previously published mass spectrometry analysis of the *Xenopus laevis* oocyte plasma membrane (Hill *et al*., 2005) to guide our investigation of bilayer composition, and identified phosphatidylethanolamine (PE) as a promising candidate phospholipid that may regulate cytokinetic signaling. PE is a major component of the *Xenopus* plasma membrane and has been shown to be required for cytokinesis in mammalian cells (Emoto and Umeda, 2000, 2001). Therefore, we asked whether SLBs containing physiological levels of PE (20%) alter cortical Rho and F-actin dynamics compared to the previously tested SLB composition. We assayed active Rho and F-actin dynamics in the artificial cortex using total internal reflection fluorescence (TIRF) microscopy and recombinant probes for active Rho (GFP-rGBD, rhotekin GTPase binding domain (Benink and Bement, 2005)) and F-actin (Alexa fluor-labeled UtrCH, Utrophin calponin homology domain)). On bilayers containing 20% PE, we observed a dramatic change in the organization of large-scale traveling waves (Figure 1A, B). In agreement with previous findings (Landino *et al*., 2021), traveling waves on control SLBs only occurred once, emerging from multiple foci within a field of view. Upon interaction, competing wave fronts extinguished one another, terminating wave propagation. On PE-containing SLBs, traveling wave fronts were more continuous, spanning the entire field of view (Figure 1A, Video S1). These wave fronts sometimes merged upon interaction (Figure 3A, see arrowheads 1 and 2), and frequently showed more than one round of wave progression (Figure 3B-C, see arrowheads). We generally observed that PE-containing bilayers supported two or three rounds of traveling wave formation before transitioning to standing oscillations. This result demonstrates that different SLB compositions are sufficient to alter cortical patterning *in vitro*.

**Figure 1:**
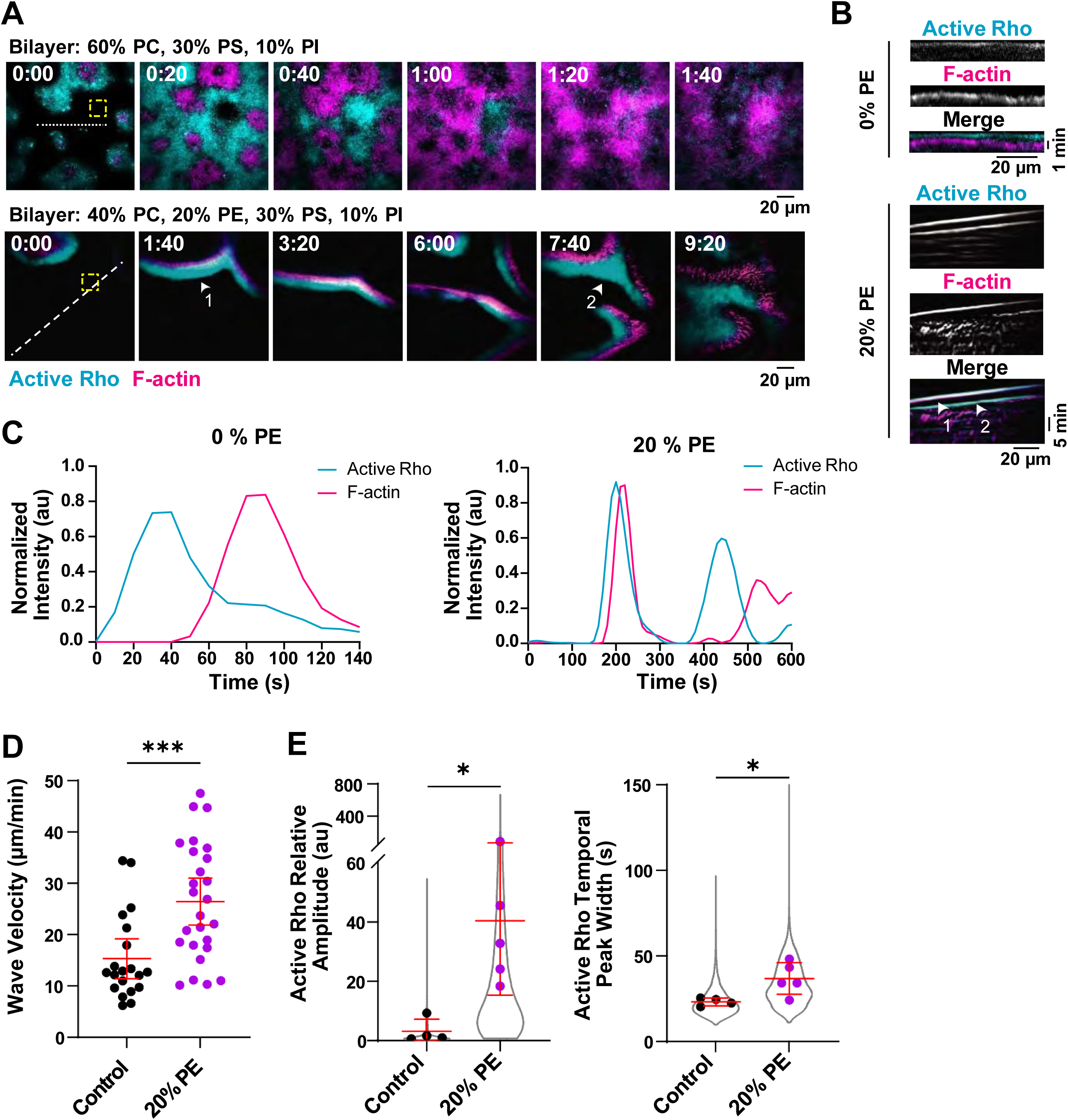
PE-containing SLBs alter active Rho and F-actin traveling wave dynamics. A) Representative micrographs of traveling waves on SLBs containing 0% or 20% PE. Arrowheads indicate merging wave fronts. Dashed lines indicate regions used to generate kymographs in (B). Yellow dashed boxes indicate regions used to measure representative intensity traces in (C). Time is indicated in minutes:seconds after the start of imaging. B) Kymographs of active Rho and F-actin generated from the dashed lines shown in (A) for traveling waves in LSS on 0% or 20% PE-containing SLBs. Arrowheads indicate consecutive wave fronts. C) Representative intensity over time traces of active Rho and F-actin traveling waves on 0 and 20% PE-containing SLBs. Generated from the regions of interest (yellow dashed boxes) shown in (A). D) Velocity of traveling waves on control (0% PE) or 20% PE SLBs. Red bars indicate mean ± 95% CI. For control, n = 20 wave fronts from 4 videos from 3 extract preps on 3 experimental days. For 20% PE, n = 26 wave fronts from 5 videos from 4 extract preps on 4 experimental days, *** p ≤ 0.001. E) Quantification of relative amplitude and temporal width of active Rho traveling waves on control (0%, black) and 20% PE-containing SLBs (magenta). Data points from all experiments are represented by a violin plot (gray) and the mean of each experiment is shown as a dot. Red bars indicate total mean ± SD of the experimental means. For control, n = 3083 values from 4 videos from 3 biological samples from 3 experimental days. For 20% PE, n = 5288 values from 5 videos from 5 biological samples from 4 experimental days. * p ≤ 0.05.

We quantified traveling wave velocity using kymographs to track the distance of wave front movement over time and found that traveling waves moved faster on SLBs containing PE (Figure 1D). We also quantified temporal wave dynamics using high-throughput automated wave analysis code (Landino *et al*., 2021; Swider *et al*., 2022) to determine the temporal shift between active Rho and F-actin peaks, wave amplitude and temporal peak width. Traveling waves showed a significant increase in the relative amplitude of active Rho, and a corresponding larger temporal width (Figure 1E). We observed no change in the temporal shift between active Rho and F-actin peaks in traveling wave (Figure S1A). These results suggest that SLBs containing PE support increased active Rho positive feedback in the artificial cortex.

Given that phospholipids are sometimes patterned alongside cortical F-actin waves in cells (Tong *et al*., 2023), we also asked whether PE was patterned within traveling waves. To test this, we used Rhodamine-PE to directly image phospholipid localization within PE-containing SLBs. We found that Rhodamine-PE was not patterned in a manner that correlated to traveling waves of active Rho or F-actin (Figure S1C-D). This result agrees with previous findings that Cy5-PC is not patterned within the SLB (Landino *et al*., 2021), and indicates that these phospholipids are evenly distributed within the bilayer and do not organize in a manner that correlates to active Rho or F-actin localization.

Our previous characterization of the artificial cortex found that the appearance of traveling waves positively correlated with more fluid bilayers (Landino *et al*., 2021). We predicted that PE-containing SLBs might be more fluid than control compositions, and therefore able to support additional rounds of traveling wave formation. We assayed the fluidity of PE-containing SLBs by fluorescence recovery after photobleaching (FRAP). Surprisingly, we found that Cy5-labeled PC in bilayers containing PE, was less fluid after photobleaching as measured by both the mobile fraction (percent recovery) and the half time (t_1/2_) across a range of PE concentrations (0%, 10%, 20%, or 30% PE, Figure S1E-F). Similar results were obtained using Rhodamine-PE (Figure S1E-F), suggesting that overall SLB fluidity is reduced when PE is present. This result indicates that the addition of PE to the bilayer does not alter traveling or oscillatory dynamics by increasing the fluidity of the SLB, and the effect of PE on cortical patterning is likely to occur through another mechanism.

### PE-containing SLBs alter standing active Rho and F-actin oscillations

After traveling waves end, active Rho and F-actin dynamics in the artificial transition to standing oscillations (Landino *et al*., 2021). We observed that both control and 20% PE-containing SLBs support standing Rho and F-actin oscillations (Figure 2A-C, Video S2). We quantified oscillations on SLBs with or without 20% PE and found that active Rho and F-actin relative amplitude increased on PE-containing SLBs, as well as F-actin temporal width (Figure 2D). Like with traveling waves, we observed no significant change in the temporal shift between active Rho and F-actin peaks (Figure S1B), suggesting that the addition of PE does not alter the time between active Rho signaling and actin assembly. These results indicate that the presence of PE in the bilayer alters the dynamics of active Rho and F-actin standing oscillations, and support a model where PE amplifies positive feedback signaling.

**Figure 2:**
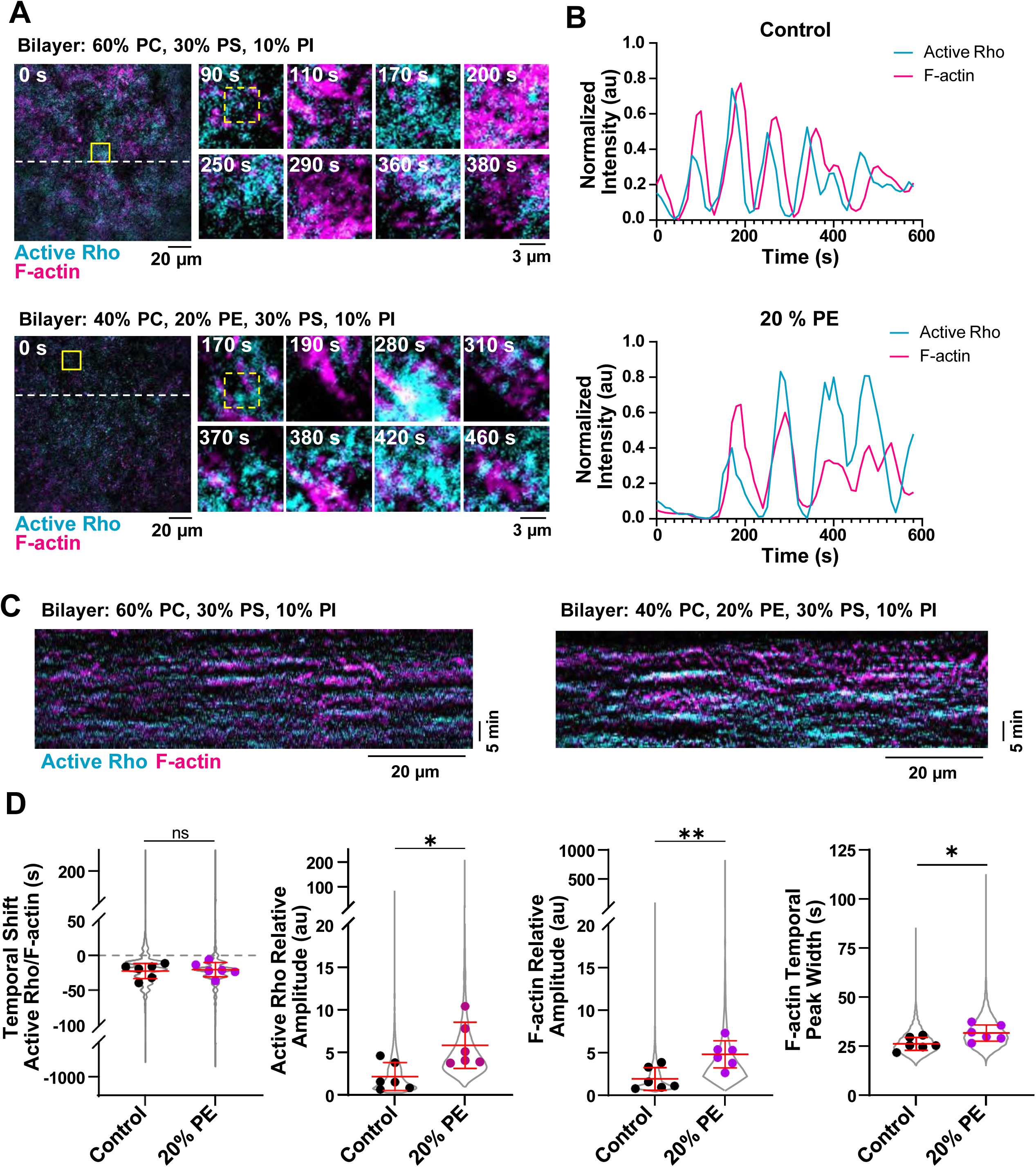
PE-containing SLBs alter the dynamics of active Rho and F-actin standing oscillations. A) Representative micrographs of standing oscillations on SLBs containing 0% or 20% PE. Time is shown in seconds (s). Left: micrograph of the whole field of view at time = 0 s. Dashed lines indicate regions used to generate kymographs in (C). Yellow solid boxes indicate regions of interest used to generate enlarged micrographs. Right: Enlarged micrographs of standing oscillations generated from the regions indicated by the solid yellow box. Dashed yellow box indicates region used to generate representative intensity traces shown in (B). B) Representative intensity over time traces of active Rho and F-actin standing oscillations on 0 and 20% PE-containing SLBs. Generated from the regions of interest (yellow dashed boxes) shown in (A). C) Representative kymographs of standing oscillations on control (0%) or 20% PE-containing SLBs. D) Quantification of the temporal shift, active Rho and F-actin relative amplitude, and F-actin temporal peak width in standing oscillations in LSS on control (0% PE, black) and 20% PE-containing (magenta) SLBs. Values from all experiments are represented by the violin plot (gray) and the mean of each experiment is shown as a dot. Red bars indicate total mean ± SD of the experimental means. For control, n = 7415 values from 6 videos from 5 biological samples from 4 experimental days. For 20% PE, n = 9754 values from 6 videos from 4 biological samples from 2 experimental days. * p ≤ 0.05, ** p ≤ 0.01.

**Figure 3:**
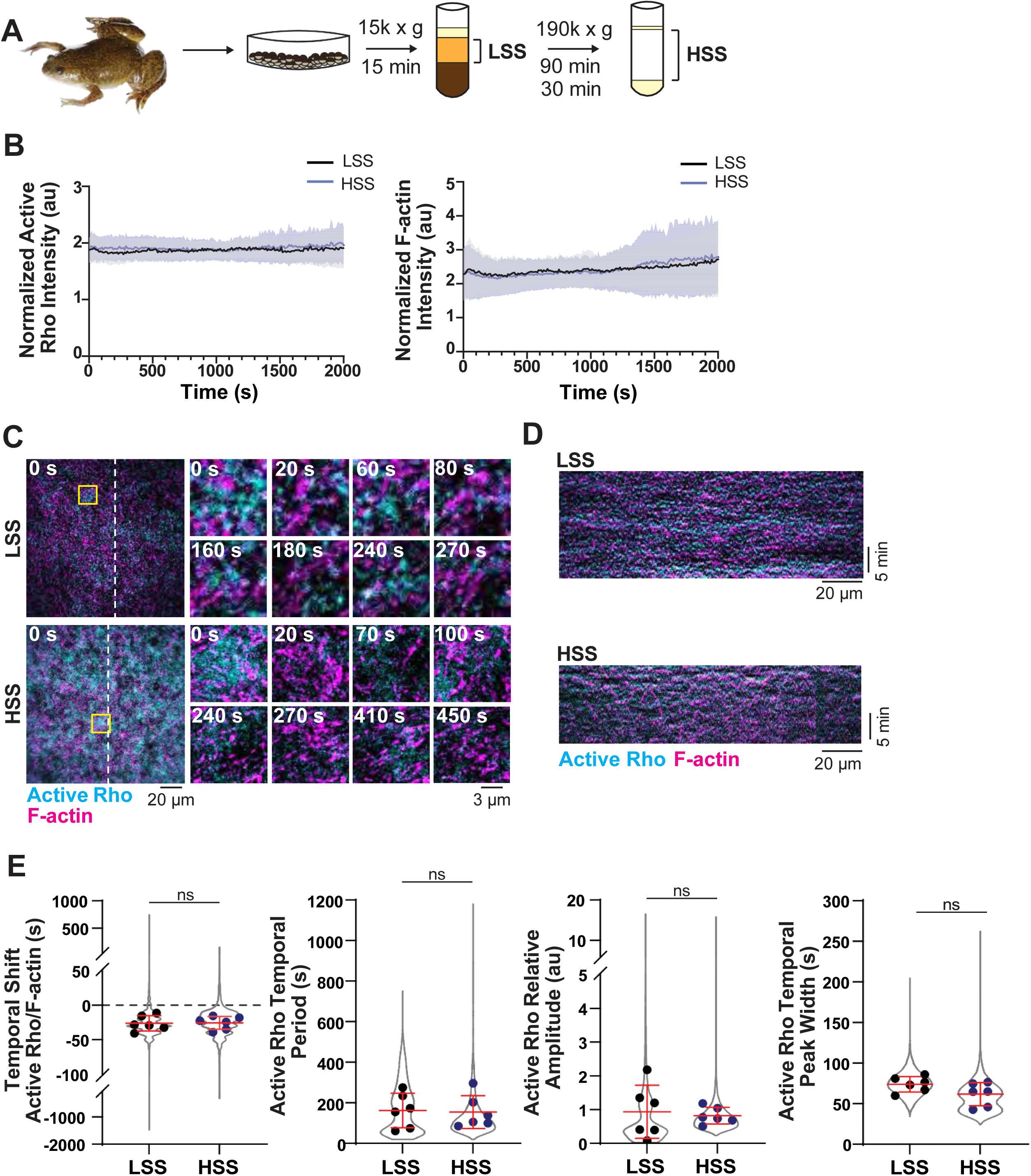
Active Rho and F-actin self-organize in the artificial cortex independent of intracellular organelles. A) Schematic of low-speed supernatant (LSS) or high-speed supernatant (HSS) preparation for use in the artificial cell cortex. Time indicates length of centrifugation; 15 min to generate LSS, and subsequent 90 and 30 min centrifugation to generate HSS. B) Normalized whole-field intensity of active Rho and F-actin over time in LSS (black) and HSS (blue) added to supported lipid bilayers. Mean (solid line) ± standard deviation (SD, shading) is shown. For LSS, n = 7 videos from 4 extract preps from 3 experimental days, for HSS, n = 13 movies from 9 extract preps from 6 experimental days. C) Representative micrographs of reconstituted active Rho and F-actin standing oscillations in the artificial cortex in LSS or HSS. Dashed lines indicate regions used to generate kymographs in (D). Left: whole field of view at time = 0. Yellow box indicates region used to generate enlarged images on the right. Right: enlarged micrographs showing active Rho and F-actin standing oscillations in LSS or HSS. Time is indicated in seconds after the start of oscillatory dynamics. D) Kymographs of active Rho and F-actin oscillations over time in LSS or HSS. Generated from the corresponding dashed lines in (C). E) Quantification of oscillatory dynamics in LSS (black) or HSS (blue) including temporal shift, active Rho temporal period, relative amplitude and temporal peak width. Values from all experiments are represented by the violin plot (gray) and the mean of each experiment is shown as a dot. Red bars indicate total mean ± standard deviation (SD) of the experimental means. For LSS, n = 6352 values from 6 videos from 4 biological samples from 3 experimental days, For HSS, n = 4753 values from 6 videos from 4 biological samples from 3 experimental days.

### Self-organized active Rho and F-actin oscillations do not require intracellular organelles

Our investigation of active Rho and F-actin oscillations in the artificial cortex system thus far used low-speed centrifugation to isolate actin-intact cytoplasmic extract (low-speed supernatant, LSS) from *Xenopus laevis* eggs. This LSS fraction lacks plasma membrane but contains a variety of intracellular membrane-bound organelles (Field *et al*., 2014; Landino *et al*., 2021). We considered that cytoplasmic lipids from LSS could directly interfere with the composition of the SLB in the artificial cortex. Moreover, experiments in the *Xenopus laevis* embryo indicate that local calcium signaling can impact cortical excitability, suggesting a possible role for cortically associated endoplasmic reticulum in regulating F-actin dynamics (Sepaniac *et al*., 2023). To test whether cytosolic membrane-bound organelles were required for reconstitution of self-organized cortical dynamics, we performed high-speed centrifugation (high-speed supernatant, HSS) to deplete intracellular organelles from our cytoplasm preparation (Figure 3A, (Sheehan *et al*., 1988; Hoogenboom *et al*., 2017)). We validated the reduction of cytoplasmic phospholipids in HSS using thin layer chromatography and lipidomics (Figure S2A-C). The major detectable lipids in the LSS were reduced to undetectable levels by TLC analysis (Figure S2A) and all major lipid class identified by lipidomic analysis were significantly reduced (Figure S2B-C) including mitochondrially-enriched cardiolipin (Paradies *et al*., 2019). This indicates that high-speed centrifugation did indeed remove membrane-bound organelles.

We next assayed LSS and HSS for active Rho and F-actin intensity and dynamics on control SLB compositions (60% PC, 30% PS, 10% PI). We first observed that total active Rho and F-actin intensities on the bilayer were similarly maintained over 60 minutes in both LSS and HSS (Figure 3B). We found that HSS supported both traveling waves (Figure S3A) and standing oscillations, however, standing oscillations are more consistently repeatable between experiments. Therefore, we focused on quantifying active Rho and F-actin oscillatory dynamics for the rest of our studies (Figure 3C-D, S2D, Video S3). We found no significant difference in the temporal shift between active Rho and F-actin peaks, active Rho or F-actin temporal periods, relative amplitudes, or temporal peak widths in HSS compared to LSS (Figure 3E, Figure S2E). This finding demonstrates that intracellular, membrane-bound organelles are not required for self-organized Rho and F-actin oscillations. Additionally, the reduction of cytoplasmic lipids in HSS reduces the possibility that cytoplasmic lipids will interfere with bilayer fluidity or composition in our studies.

### PE-containing SLBs alter standing oscillations in HSS

Having established HSS as an experimental tool for studying self-organized active Rho and F-actin oscillations, we next investigated the dynamics of standing oscillations in HSS on SLBs containing a range of PE concentrations (0%, 10%, 20%, or 30% PE, Figure 4A-C). While HSS supported traveling wave formation on SLBs with and without PE (Figure S3A), we focused on quantifying the more reproducible oscillatory dynamics. We observed that an increasing fraction of PE in the SLBs caused increased relative amplitude of active Rho (10% PE) and F-actin (30% PE, Figure 4C, Video S4). We also observed an increased F-actin temporal period (30% PE, Figure 4C). Notably, increasing the fraction of PE within SLB correlated with a greater degree of variability between experiments. In agreement with our findings from traveling waves and standing oscillations in LSS (Figure S1A-B), we did not observe a change in the temporal shift between the active Rho and F-actin peaks (Figure S3), indicating that the addition of PE does not alter the kinetics of active Rho signaling to F-actin assembly. Taken together with our studies in LSS, our findings show that PE-containing SLBs support higher amplitude traveling waves and standing oscillations and a longer temporal F-actin period, without altering the relationship between active Rho and F-actin.

**Figure 4:**
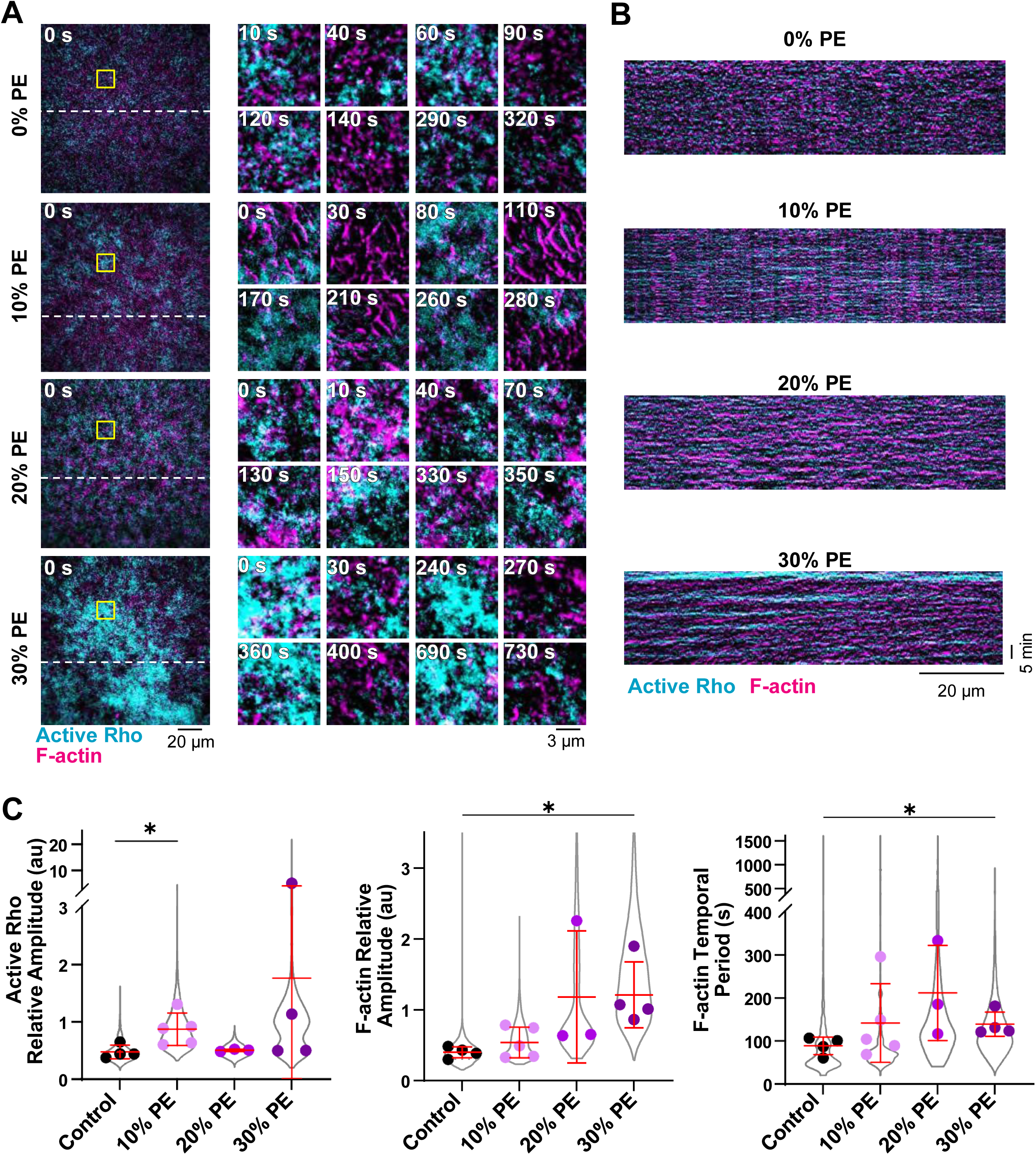
Active Rho and F-actin amplitude of cortical oscillations increases in HSS on PE-containing SLBs. A) Representative micrographs of standing oscillations on SLBs containing 0%, 10%, 20% or 30% PE. Time is shown in seconds (s). Left: micrographs of the whole field of view at time = 0 s. Dashed lines indicate regions used to generate kymographs in (B). Yellow solid boxes indicate regions of interest used to generate enlarged micrographs. Right: enlarged micrographs of standing oscillations generated from the regions indicated by the solid yellow box. B) Representative kymographs of active Rho and F-actin standing oscillations in HSS on 0%, 10%, 20% or 30% PE-containing SLBs. C) Quantification of active Rho and F-actin relative amplitude, and F-actin temporal period in HSS on SLBs containing 0% (control, black), 10% (lavender), 20% (magenta), or 30% (purple) PE. Values from all experiments are represented by the violin plot (gray) and the mean of each experiment is shown as a dot. Red bars indicate total mean ± SD of the experimental means. For control, n = 3872 values from 4 videos from 3 biological samples from 3 experimental days. For 10% PE, n = 5099 values from 5 videos from 3 biological samples from 3 experimental days. For 20% PE, n = 3354 values from 3 videos from 3 biological samples from 2 experimental days. For 30% PE, n = 8222 values from 4 videos from 4 biological samples from 3 experimental days. * p ≤ 0.05.

### PIP_2_ supports cortical F-actin assembly

We next used HSS to investigate whether PIP_2_ within the membrane alters active Rho and F-actin dynamics. PIP_2_ has been identified as a key regulatory lipid, and is thought to support Rho signaling, F-actin assembly, and cytokinesis (Field *et al*., 2005; Liu *et al*., 2012; Janmey *et al*., 2018; Budnar *et al*., 2019). We asked whether bilayers with or without PIP_2_ (0.1%, (Landino *et al*., 2021)) similarly supported cortical F-actin assembly and active Rho and F-actin oscillations. We used HSS to avoid possible contamination of our SLB composition by cytoplasmic lipids found in LSS. We first tested whether F-actin assembled on SLBs with or without PIP_2_ and found that the presence of 0.1% PIP_2_ dramatically increased the propensity for F-actin assembly on the SLB (Figure 5A) 10 minutes after HSS addition (Figure S4A-B). This result demonstrates that PIP_2_ supports the initial phase of cortical F-actin self-assembly. As our SLBs contain PI, a substrate for PIP_2_ synthesis by PI4-Kinase (PI4K) (Czech, 2000), we also considered that conversion of PI to PIP_2_ could explain the fraction of SLBs (50%) that support F-actin assembly without PIP_2_. We tested this possibility by pre-treating HSS with a selective inhibitor of PI4KIIIα (GSK-A1) to block PIP_2_ synthesis (Kilwein *et al*., 2025). We observed a sustained decrease in F-actin intensity on the SLB when HSS was treated with GSK-A1 compared to DMSO-treated controls (Figure 5B). This finding suggests that F-actin assembly on bilayers lacking PIP_2_ is supported by conversion of PI into PIP_2_ by PI4K in our system.

**Figure 5:**
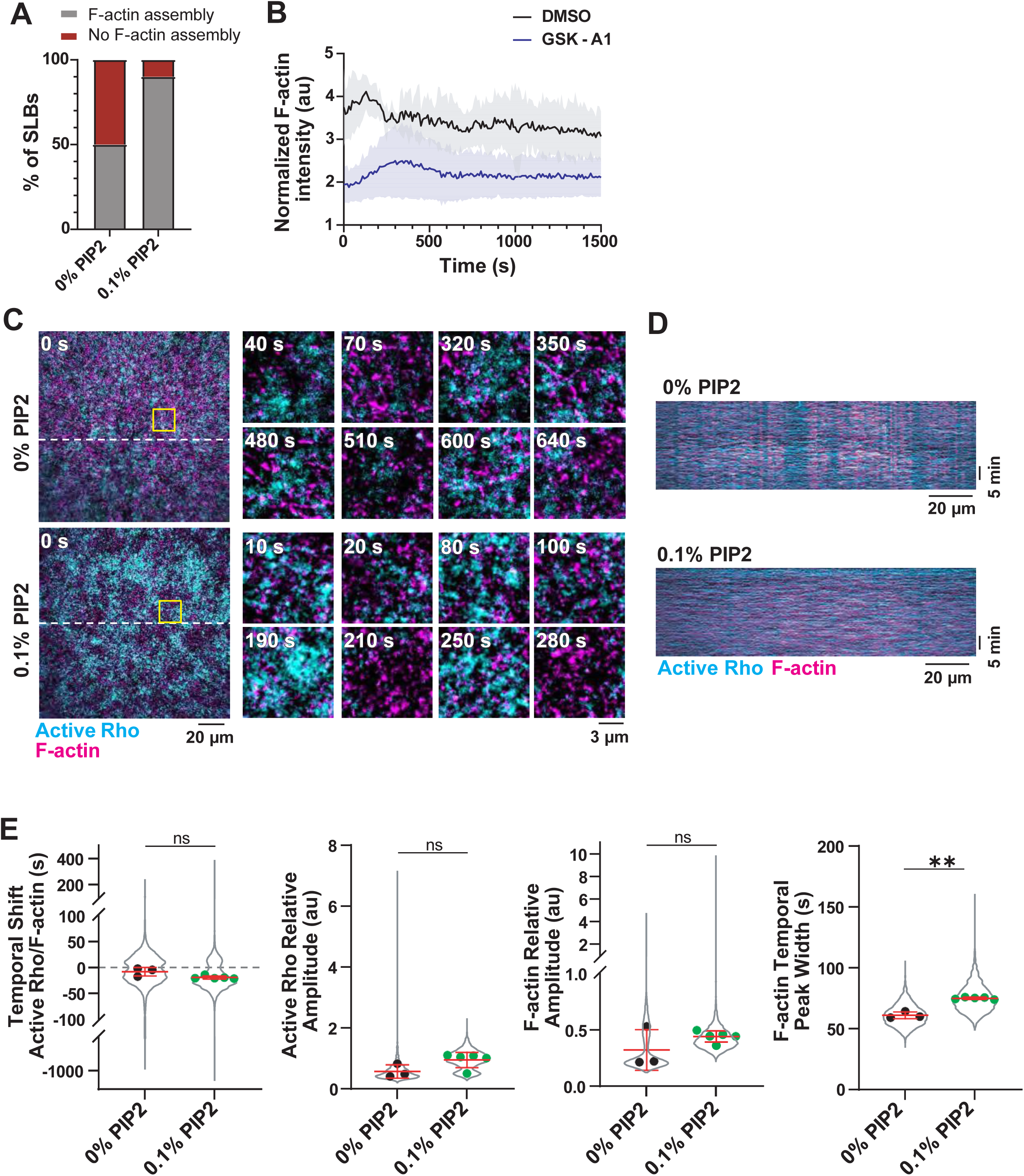
PIP_2_ supports F-actin assembly on SLBs. A) Quantification F-actin assembly on SLBs containing 0% or 0.1% PIP_2_ after 10 minutes. n = 20 paired movies from 10 biological replicates. B) Quantification of normalized F-actin intensity over time after treatment with DMSO or PI4KIIIɑ inhibitor (GSK-A1). Mean (solid line) ) ± standard deviation (SD, shading) is shown. For both n = 3 videos from 3 extract preps on 2 experimental days. C) Representative micrographs of oscillatory dynamics in HSS on SLBs containing 0 or 0.1% PIP_2_. Left: micrographs of the whole field of view at time = 0 s. Dashed lines indicate regions used to generate kymographs in (D). Time is indicated in seconds after the start of oscillatory dynamics. Yellow solid boxes indicate regions of interest used to generate enlarged micrographs. Right: enlarged micrographs of standing oscillations generated from the regions indicated by the solid yellow box. D) Kymographs of active Rho and F-actin oscillations over time in LSS or HSS. Generated from the corresponding dashed lines in (C). E) Quantification of oscillatory dynamics in HSS on SLBs containing 0% (black) or 0.1% (green) PIP_2_. Values from all experiments are represented by the violin plot (gray) and the mean of each experiment is shown as a dot. Red bars indicate total mean ± SD of the experimental means. For 0% PIP_2_ SLBs, n = 3453 values from 3 videos from 3 biological samples from 2 experimental days. For 0.1% PIP_2_ SLBs, n = 5618 values from 5 videos from 3 biological samples on 2 experimental days.

We then assayed dynamics of active Rho and F-actin on SLBs with or without 0.1% PIP_2_. For this analysis, we only compared experiments where we observed robust F-actin assembly on the SLB (50%, Figure 5A). We found few differences in cortical oscillations over time on SLBs with or without 0.1% PIP_2_ (Figure 5C-E, S4C, Video S5), although we did observe moderate but statistically insignificant increases for the relative amplitude of active Rho and F-actin peaks on SLBs with 0.1% PIP_2_ (Figure 5E). We also observed an increased F-actin temporal width on bilayers with 0.1% PIP_2_ (Figure 5E).These results show that the initial presence of PIP_2_ in the SLB supports robust F-actin assembly and establishment of a cortical actin network, and that PIP_2_ is likely replenished within the SLB by phosphoinositide metabolism. Once cortical oscillations are established in the artificial cortex, they are overall similar on bilayers with and without PIP_2_.

We also considered that bilayers with and without PIP_2_ may have different membrane fluidity and we tested this possibility using FRAP of Cy5-labeled PC. We observed no significant difference in recovery of bilayers with or without PIP_2_, indicating that PIP_2_ does not meaningfully alter bilayer fluidity (Figure S4D).

### Sphingomyelin and cholesterol-containing bilayers alter cortical oscillation dynamics

Our finding that traveling waves and standing oscillations of active Rho and F-actin change on bilayers containing PE lead us to predict that overall bilayer composition is important for determining self-organized cortical dynamics. We next investigated oscillatory dynamics on SLBs containing lipids previously identified as major components of the *Xenopus* oocyte plasma membrane, including cholesterol (20% of total membrane mass) and sphingomyelin (25% of phospholipid mass) (Hill *et al*., 2005). Previous studies suggest cholesterol and sphingomyelin localize to the cytokinetic furrow and support successful cytokinesis (Fernandez *et al*., 2004; Ng *et al*., 2005; Abe *et al*., 2012), however whether they regulate active Rho and F-actin oscillations has not been tested. We investigated this possibility using SLB compositions containing physiological levels of cholesterol or sphingomyelin (Figure 6A, hereafter referred to as “cholesterol” or “sphingomyelin” SLBs respectively, (Hill *et al*., 2005)).

**Figure 6:**
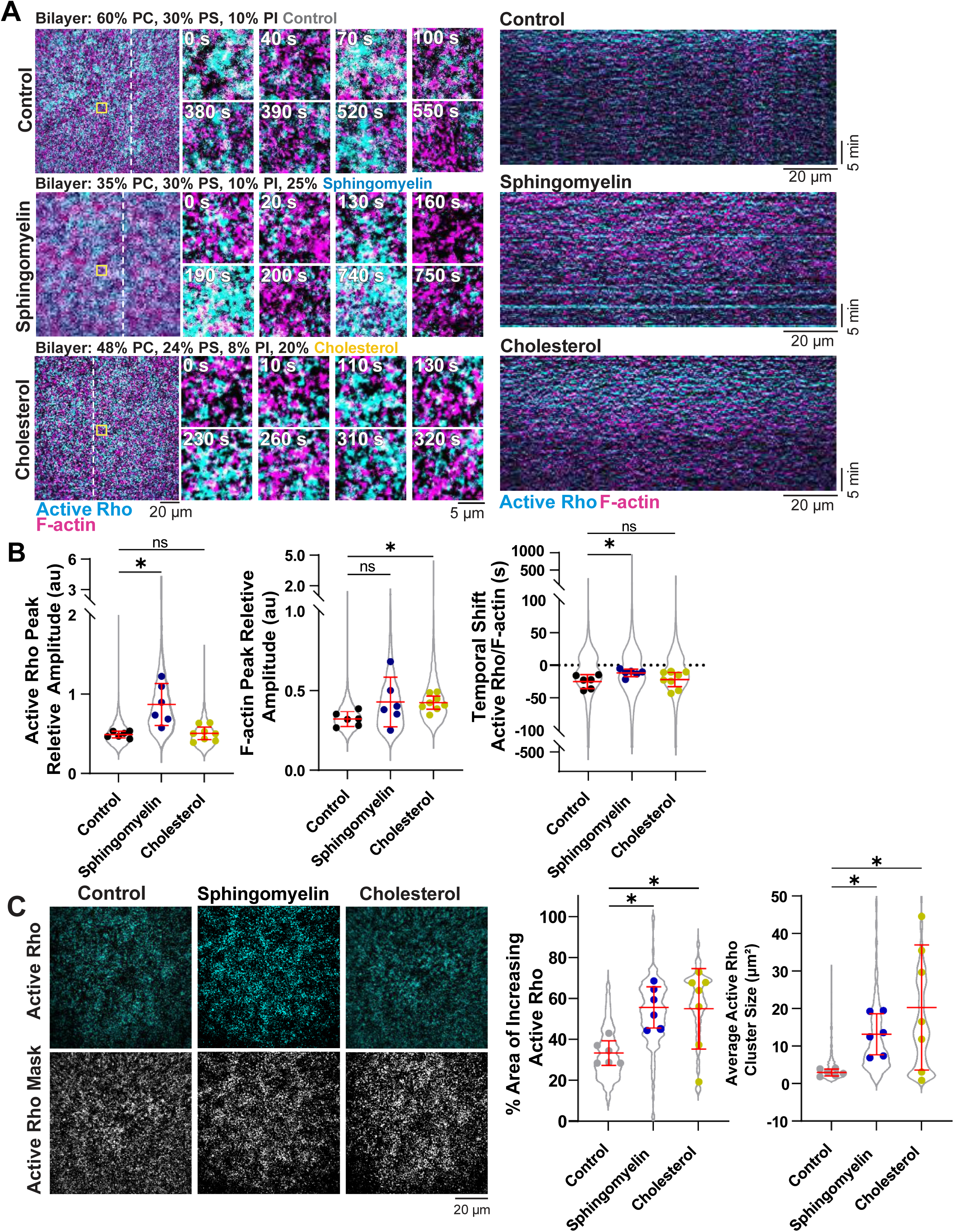
Temporal active Rho and F-actin oscillations are altered on sphingomyelin and cholesterol-containing bilayers. A) Left: Representative micrographs of active Rho (cyan) and F-actin (magenta) oscillations on control, sphingomyelin, and cholesterol-containing bilayers. Time is indicated in minutes after the onset of oscillatory dynamics. The white dotted line indicates the region used to create kymographs. Right: kymographs generated from the dashed lines of active Rho (cyan) and F-actin (magenta) oscillations on control, sphingomyelin, and cholesterol-containing bilayers. B) Quantification of the active Rho and F-actin peak relative amplitude and active Rho/F-actin temporal shift in HSS on control, sphingomyelin, and cholesterol-containing SLBs. All data points are represented in the violin plot and the average for each experiment is shown as a single dot. Red bars indicated the total average and standard deviation. For controls, n = 5623 values from 6 videos from 3 biological replicates across 4 experimental days. For sphingomyelin-containing SLBs, n = 6606 values from 6 videos from 3 biological replicates across 4 experimental days. For cholesterol-containing SLBs, n = 6647 values from 8 videos from 6 biological replicates across 3 experimental days. * p ≤ 0.05. C) Left: Representative images of the active Rho intensity on control, cholesterol and sphingomyelin SLBs and representative thresholded masks used to calculate the percent area of increasing Rho activity across time. Right: Quantification of percent area of increased active Rho at each frame in the video. Data points in the violin plot represent each frame of all videos and the average for each video is shown as a single dot. Red bars indicated the average of the experimental means and standard deviation. For controls, n = 1000 values from 6 videos from 3 biological replicates across 4 experimental days. For cholesterol-containing SLBs, n = 991 values from 8 videos from 6 biological replicates across 3 experimental days. For sphingomyelin-containing SLBs n = 868 values from 6 videos from 3 biological replicates across 4 experimental days. * p ≤ .05.

We considered that the presence of cholesterol or sphingomyelin might alter SLB fluidity, so we first characterized the dynamics of cholesterol and sphingomyelin SLBs (Cy5-labed PC) using FRAP. We found no discernable difference between the control and cholesterol nor the control and sphingomyelin bilayers, as assayed by the recovery rate (t_1/2_) or mobile fraction (percent recovery, Figure S5A). These results show that the membrane fluidity of bilayers containing cholesterol or sphingomyelin is unchanged in our system.

We next investigated active Rho and F-actin temporal dynamics of active Rho and F-actin standing oscillations on cholesterol and sphingomyelin-containing bilayers using HSS. To increase the robustness of F-actin polymerization (Figure 2B), we added 0.1% PIP_2_ to each bilayer composition. Control, cholesterol, and sphingomyelin SLBs all produced active Rho and F-actin standing oscillations over time (Figure 6A, Video S6). We quantified oscillatory dynamics of active Rho and F-actin on all three bilayer compositions and found that the active Rho relative amplitude increased on sphingomyelin SLBs compared to control SLBs (Figure 6B). We also found a significant change in the F-actin relative amplitude for the cholesterol bilayers (Figure 6B). We did not observe changes in the period, temporal width, on cholesterol or sphingomyelin-containing SLBs compared to controls (Figure S4B). We also observed that the temporal shift in the active Rho and F-actin peaks was reduced on sphingomyelin bilayers, suggesting faster signaling between Rho activity and F-actin assembly (Figure 6B). The temporal shift was unchanged on cholesterol bilayers (Figure 6B). The increased relative amplitude of active Rho and F-actin, on sphingomyelin and cholesterol SLBs respectively, suggests that there is increased positive feedback signaling and F-actin assembly on these bilayers compared to controls. These results show that SLB composition modulates active Rho and F-actin signaling within the artificial cortex.

We also observed differences in the spatial organization of active Rho on cholesterol and sphingomyelin SLBs. We quantified this by measuring the total area of increasing active Rho signal on control, cholesterol, and sphingomyelin SLBs with a mask of the active Rho intensity throughout the video (Figure 6C) (Swider *et al*., 2022). We measured the total area of the mask at each time point to determine the percent area Rho activity in the field of view on each SLB composition. We also measured the average size of individual clusters of active Rho using this method. We found that there was a significant increase in the percent area of dynamic active Rho on cholesterol and sphingomyelin SLBs (Figure 6C). We also found that the size of individual active Rho clusters was larger on cholesterol and sphingomyelin SLBs (Figure 6C). This result suggests that membrane composition is important for spatial activation of Rho, in addition to regulating temporal dynamics.

### Rate of active Rho binding and F-actin assembly is reduced on cholesterol and sphingomyelin bilayers

During our characterization of active Rho and F-actin standing oscillations on sphingomyelin and cholesterol SLBs, we also observed a delay in the initial cortical F-actin polymerization on sphingomyelin or cholesterol SLBs. We hypothesize that the presence of sphingomyelin or cholesterol in the bilayer could alter the rate of cortical assembly. To precisely test this, we performed live imaging of extract addition to the SLB, with shorter sampling intervals (2 seconds) than we previously used to image cortical oscillations (10 seconds). On control SLBs, we observed that there was a noticeable increase in active Rho signal on the bilayer before the onset of robust actin polymerization. It took approximately 40 seconds from the appearance of active Rho for detectable actin polymerization to begin (Figure 7A-B, Video S7), a timeframe that closely aligns with the reported temporal shift between active Rho and F-actin oscillations *in vivo* and *in vitro* (Bement *et al*., 2015; Landino *et al*., 2021). On sphingomyelin and cholesterol SLBs, there was a delay in the time from extract addition to active Rho signal and robust actin polymerization compared to control SLBs (Figure 7C-D). This delayed onset correlated to an increased temporal shift between active Rho and F-actin on sphingomyelin and cholesterol bilayers (Figure 7D). These results demonstrate that bilayer composition regulates the kinetics of Rho activation on and F-actin assembly on the bilayer during the initial establishment of the reconstituted cortex.

**Figure 7:**
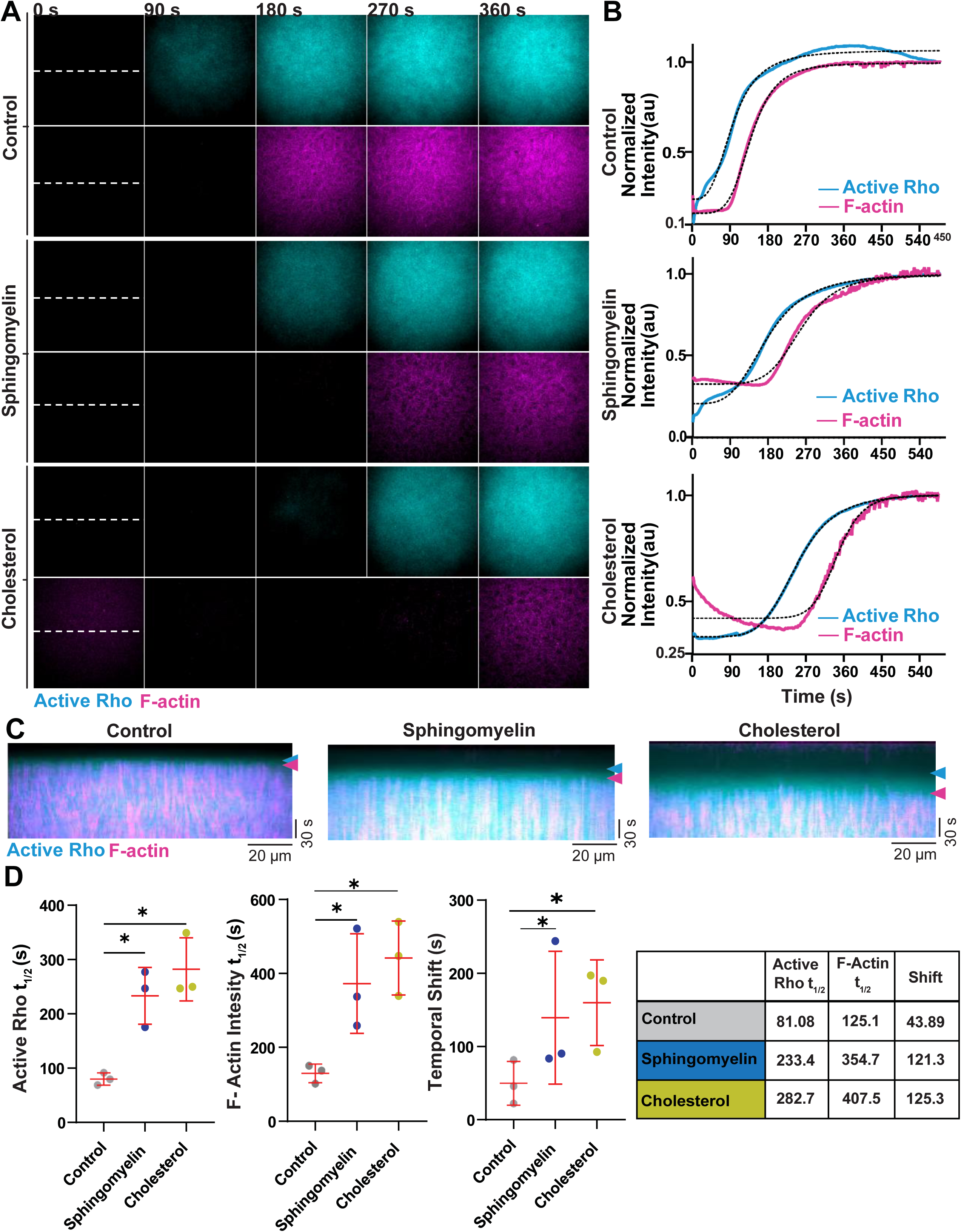
Onset of active Rho and F-actin dynamics are delayed on cholesterol or sphingomyelin-containing bilayers. A) Representative micrographs of active Rho (cyan) and F-actin (magenta) accumulation on control, sphingomyelin, and cholesterol-containing bilayers. Time is indicated in seconds after extract addition to the bilayer. The white dotted line indicates the region used to create kymographs. B) Normalized intensity curves of active Rho (cyan) and F-Actin (magenta) after extract addition to control, sphingomyelin, and cholesterol-containing bilayers. Dotted line indicates best fit sigmoidal curve. C) Kymographs of active Rho (cyan) and F-actin (magenta) generated from the dashed lines shown in (A) on control, sphingomyelin, and cholesterol-containing bilayers. Arrows indicate the onset of active Rho binding to the SLBs (cyan) and onset of detectable F-actin polymerization (magenta). D) Left: Quantification of the active Rho and F-actin SLB binding t_1/2_ and the temporal phase shift between active Rho and F-actin in HSS on control, sphingomyelin, and cholesterol-containing SLBs. Each experiment is represented as a dot and red lines indicate the mean ± SD. Right: Table of Active Rho and F-Actin t_1/2_ and temporal shift for control, sphingomyelin, and cholesterol-containing SLBs. Time is indicated in seconds. For all, n = 3 videos from 3 biological replicates on 2 experimental days. * p ≤ .05.

## DISCUSSION

In this study, we demonstrated that membrane composition alters the dynamics of active Rho and F-actin traveling waves and standing oscillations in the artificial cortex. Importantly this work represents the first reconstitution of membrane-dependent changes to self-organized active Rho and F-actin patterning. Using the artificial cortex, we have directly manipulated SLB composition to understand the role of the membrane in regulating cortical dynamics, which presents a major advantage over approaches that indirectly manipulate the plasma membrane composition in cells. Our observation that PIP_2_ supports robust assembly of cortical F-actin on SLBs and increased F-actin amplitude within oscillations agrees with previous studies in cells showing that PIP_2_ retains active Rho at the membrane (Budnar *et al*., 2019) and at the cytokinetic furrow (Field *et al*., 2005; Wong *et al*., 2005). However, in *Xenopus laevis* embryos, furrow-localized cortical oscillations have lower amplitude F-actin waves than non-furrow regions (Swider *et al*., 2022), suggesting that increased PIP_2_ at the division plane is unlikely to explain furrow-specific differences in F-actin patterning observed *in vivo*. To further understand the relationship between PIP_2_ and active Rho and F-actin dynamics, future studies will focus on investigating phosphoinositide metabolism and patterning in the context of cytokinetic waves, as has been described in other excitable systems (Hansen *et al*., 2019; Tong *et al*., 2023; Erisis and Horning, 2024).

We find that SLBs containing PE dramatically changed traveling wave organization and dynamics, inducing multiple rounds of high-amplitude waves not observed on control bilayer compositions. PE-containing bilayers also alter the dynamics of standing oscillations, which have increased active Rho and F-actin amplitude. We also found that sphingomyelin and cholesterol bilayers support increased active Rho and F-actin amplitude in standing oscillations, and that sphingomyelin supports a shortened temporal shift between active Rho and F-actin peaks (Figure 8). We propose that increased wave amplitude on our experimental SLB compositions represents increased Ect2-dependent positive feedback in the cortical excitability circuit (Figure 8, (Michaud *et al*., 2022)). This idea is also supported by the finding that the spatial activation of Rho is also more robust on sphingomyelin and cholesterol SLBs. Importantly, our data suggests that bilayer fluidity is not sufficient to explain observed changes in self-organized dynamics, as we observed little to no detectable change in the fluorescence recovery after photobleaching of the membrane compositions tested here. This work shows that the membrane composition has an important role in regulating cortical dynamics and is not limited to functioning merely as a passive platform for Rho GTPase binding and F-actin assembly.

**Figure 8:**
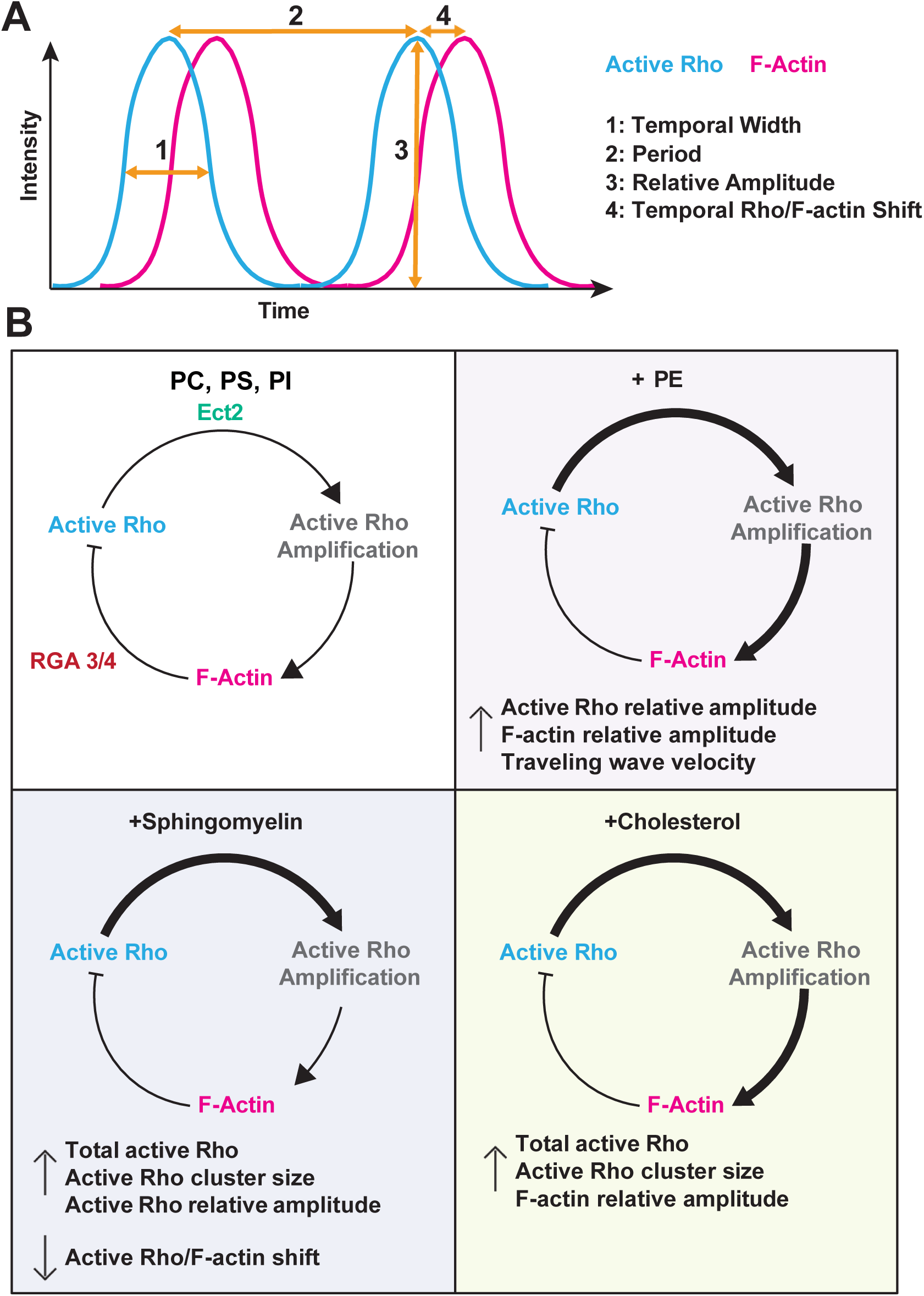
Effects of membrane composition changes to cortical oscillations schematic. A) Idealized plot of active Rho (cyan) and F-actin (magenta) cortical oscillations over time. Overlayed arrows indicate (1) temporal width, (2) period, (3) peak relative amplitude, and (4) temporal Rho/F-actin shift. B) Model of cortical excitability for control (PC,PS,PI), PE-, sphingomyelin-, and cholesterol-containing bilayers with the corresponding effect on cortical patterning. On PE SLBs, we observe increased traveling wave velocity and active Rho and F-actin amplitude suggesting increased amplification of Rho activity and increased F-actin polymerization. On sphingomyelin SLBs, we observe an increase in total active Rho, active Rho cluster size, and active Rho amplitude which suggests increased amplification of Rho activity. We also observe a decreased active Rho/F-actin temporal shift which suggests faster signaling between active Rho and F-actin assembly. On cholesterol SLBs we observe increased total active Rho, active Rho cluster size, and F-actin amplitude which suggests increased amplification of Rho activity and increased F-actin polymerization. Thicker arrows denote increased signaling and a shorter arrow indicates faster kinetics compared to controls.

We also observe that SLBs containing cholesterol and sphingomyelin showed slower Rho activation and F-actin assembly during cortex formation, which may suggest these bilayer compositions alter the kinetics of GEF-dependent activation of Rho (Armstrong *et al*., 2025). It is also possible that sphingomyelin and cholesterol indirectly regulate Rho *via* PIP_2_. Sphingomyelin is thought to regulate cytokinesis *via* PIP_2_ in cells (Emoto *et al*., 2005; Abe *et al*., 2012) and cholesterol has been suggested to regulate PIP_2_ synthesis (Rosenhouse-Dantsker, 2023). Therefore, our observed changes to cortical dynamics on sphingomyelin and cholesterol SLBs may be due to changes in PIP_2_ synthesis or organization. It is also possible that RhoA may interact directly with specific lipids. Rho is known to associate with the membrane through C-terminal prenylation, however, the protein sequence also contains a cholesterol binding motif (CARC) which could alter membrane retention times on cholesterol bilayers (Fantini and Barrantes, 2013). Deciphering the mechanistic relationship between Rho activation, diffusion, retention, and downstream signaling will be the focus of future studies.

The specific mechanism by which PE, sphingomyelin, and cholesterol alter cortical Rho and F-actin oscillations is somewhat unclear. All three groups have been shown to enrich at the cytokinetic furrow and midbody (Atilla-Gokcumen *et al*., 2014; Kunduri *et al*., 2022), although recent work suggests that there is a general increase of all lipids at the division site to support the addition of new cell surface area (Alonso-Matilla *et al*., 2024). One possibility is that these lipids alter lipid packing within the bilayer that affect RhoA binding or diffusion (Kunduri *et al*., 2022). For example, PE is thought to be cone-shaped, with acyl chains wider than the head group, which favors curved membranes (Kunduri *et al*., 2022). While our SLBs are flat, it is possible that differences in phospholipid shape (cone vs. cylinder) introduce minor defects within the bilayer that alter Rho binding or activation. Notably, we do not observe significant changes in bilayer fluidity for the compositions tested here, however it is possible that lateral movement of active Rho within the membrane could differ from lipid diffusion. Bilayer compositions that alter Rho’s ability to move laterally within the membrane could lead to changes in active Rho signal amplification and subsequent signaling for actin assembly (Figure 8). Future studies investigating reconstituted cortical patterning on curved membranes will reveal whether lipid packing defects or membrane topology directly regulate Rho activity.

## Abbreviations

F-actin: Filamentous actin
SLB: supported lipid bilayer
TIRF: Total internal reflection fluorescence
LSS: low speed supernatant
HSS: high speed supernatant
GFP: green fluorescent protein
rGBD: Rhotekin GTPase-binding domain
UtrCH: Utrophin calponin homology
FRAP: fluorescence recovery after photobleaching
PIP_2_: Phosphatidylinositol 4,5-bisphosphate
PC: phosphatidylcholine
PS: phosphatidylserine
PI: phosphatidylinositol
PE: phosphatidylethanolamine
SM: sphingomyelin

## ACKNOWLEDGEMENTS

We would like to acknowledge the members of the Landino lab for their critical feedback on this project and manuscript. We thank Ann Miller and Anthony Vecchiarelli for technical support and access to equipment for the preliminary work on this project. We thank members of the Bement lab for informing our thinking on cortical dynamics, and for sharing plasmids and recombinant proteins used in this study. We thank Dominic Chomchai for help with troubleshooting *waveanalysis* code for image analysis. The Dartmouth Cancer Center Genomics and Molecular Biology Shared Resource provided plasmid sequencing support. Recombinant protein expression was supported by Dartmouth College bioMT through NIH NIGMS grant P20-GM113132. Dalton Bioanalytics performed the mass spectrometry lipid analysis. This work was supported by NIH R00GM147826 awarded to J.L.

## SUPPLEMENTAL FIGURE LEGENDS

**Figure S1:**
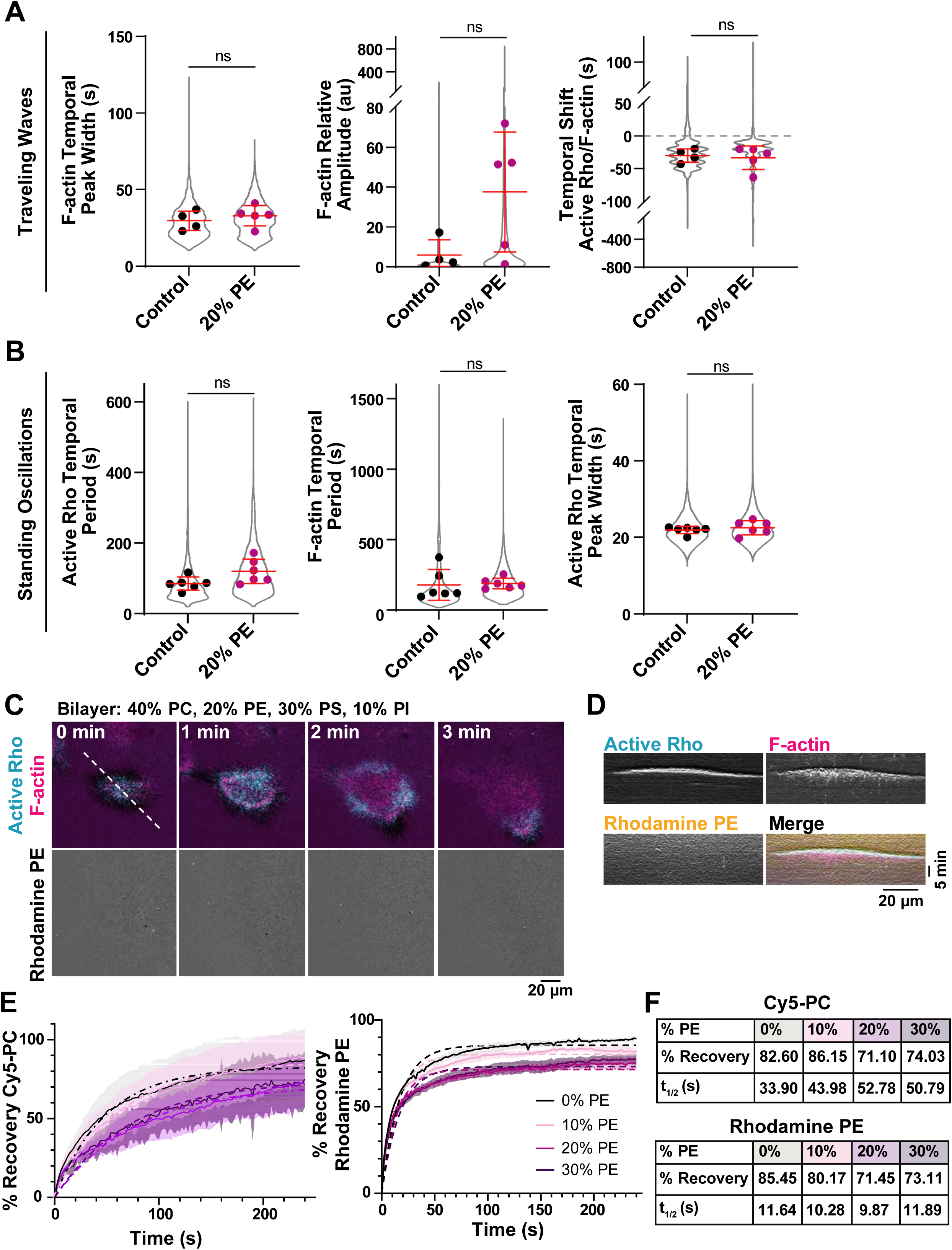
Supplemental analysis of active Rho and F-actin dynamics on PE SLBs, related to Figures 1 and 2. A) Quantification of the F-actin relative amplitude and temporal peak width and temporal shift of traveling waves on control (0%, black) or 20% PE-containing SLBs (magenta). Data points from all experiments are represented by a violin plot (gray) and the mean of each experiment is shown as a dot. Red bars indicate total mean ± SD of the experimental means. For control, n = 3083 values from 4 videos from 3 biological samples from 3 experimental days. For 20% PE, n = 5288 values from 5 videos from 5 biological samples from 4 experimental days. B) Quantification of the active Rho and F-actin temporal period and active Rho temporal width of standing oscillations in LSS on control (0%, black) or 20% PE-containing SLBs (magenta). Red bars indicate total mean ± SD of the experimental means. For control, n = 7415 values from 6 videos from 5 biological samples from 4 experimental days. For 20% PE, n = 9754 values from 6 videos from 4 biological samples from 2 experimental days. C) Representative micrographs of active Rho, F-actin, and Rhodamine PE signal in a traveling wave in LSS on 20% PE containing SLB. Time is indicated in minutes. Dashed line indicates region used to generate kymograph in (D). D) Representative kymograph of a traveling wave in LSS on a 20% PE-containing SLB, with Rhodamine-labeled PE. E) FRAP of Cy5-labeled PC or Rhodamine-labeled PE in SLBs containing 0% (black), 10% (lavender), 20% (magenta), or 30% (purple) PE. Mean (solid line) ± SD (shading) is shown. Dashed lines indicate the best fit one-phase association curve. For all, n = 6 bleach regions on 3 SLBs on 3 experimental days. F) Summary tables of % recovery and t1/2 for FRAP of Cy5-PC or Rhodamine PE as determined by one-phase association fit for recovery curves.

**Figure S2:**
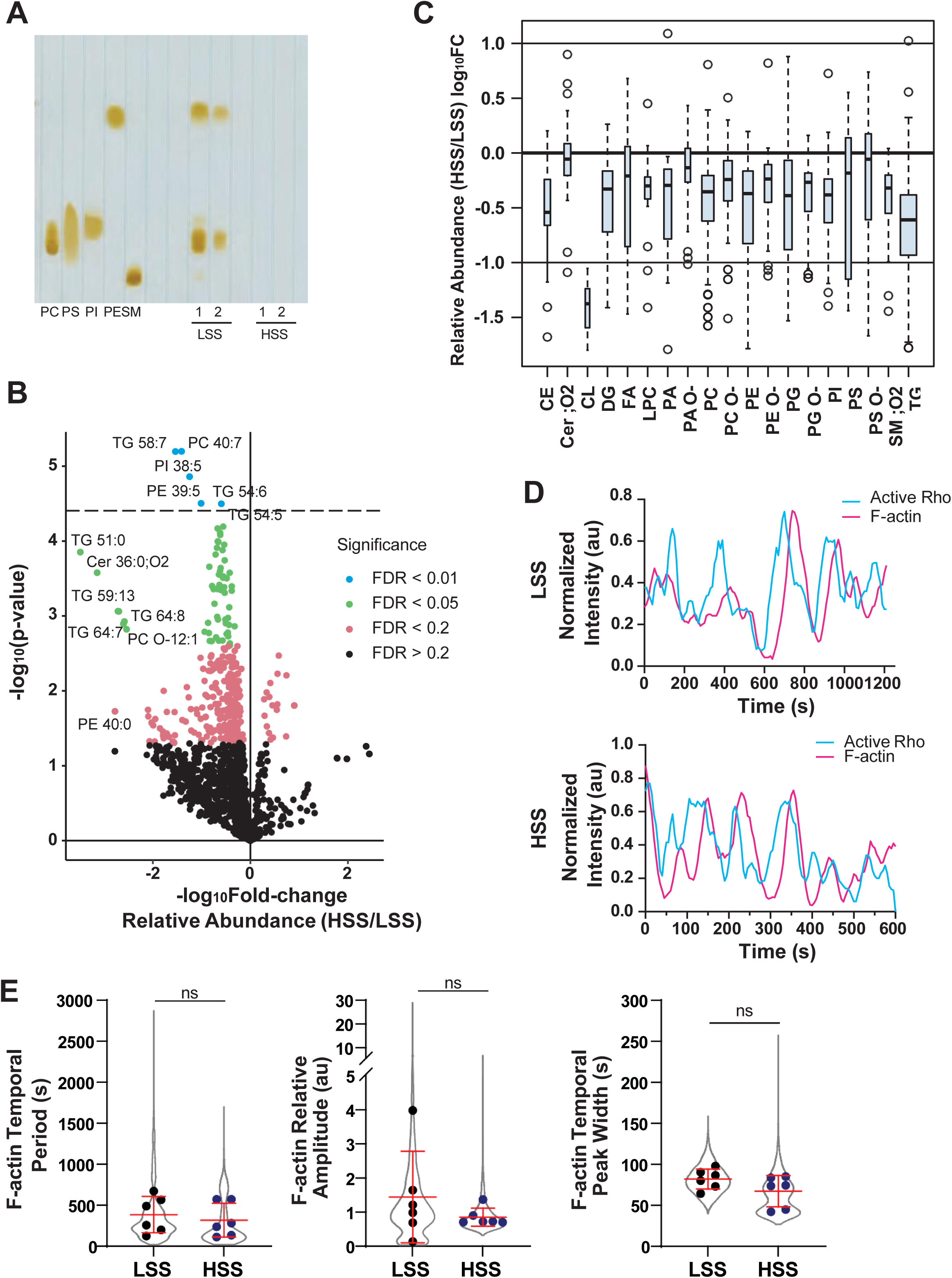
Supplemental analysis of LSS and HSS, related to Figure 3. A) Representative thin-layer chromatography (TLC) of phospholipid standards and two biological replicates of LSS or HSS cytoplasm. B) Volcano plot of lipid mass spec analysis showing the fold change of lipid abundance in HSS compared to LSS. Color coding indicates significance (false discovery rate, FDR). C) Quantification of the relative abundance of defined lipid classes in HSS compared to LSS. D) Representative plots of normalized intensity over time of active Rho and F-actin in a single 4 µm x 4 µm box from LSS and HSS. E) Quantification of F-actin standing oscillations in LSS (black) or HSS (blue) including temporal period, relative amplitude and temporal peak width. Data points from all experiments are represented by a violin plot (gray) and the mean of each experiment is shown as a dot. Red bars show the total mean ± standard deviation (SD) of the experimental means. For LSS, n = 4761 data points per video, 7 videos from 4 extract preps from 3 experimental days, For HSS, n = 4761 data points per video, 7 movies from 4 extract preps from 3 experimental days.

**Figure S3:**
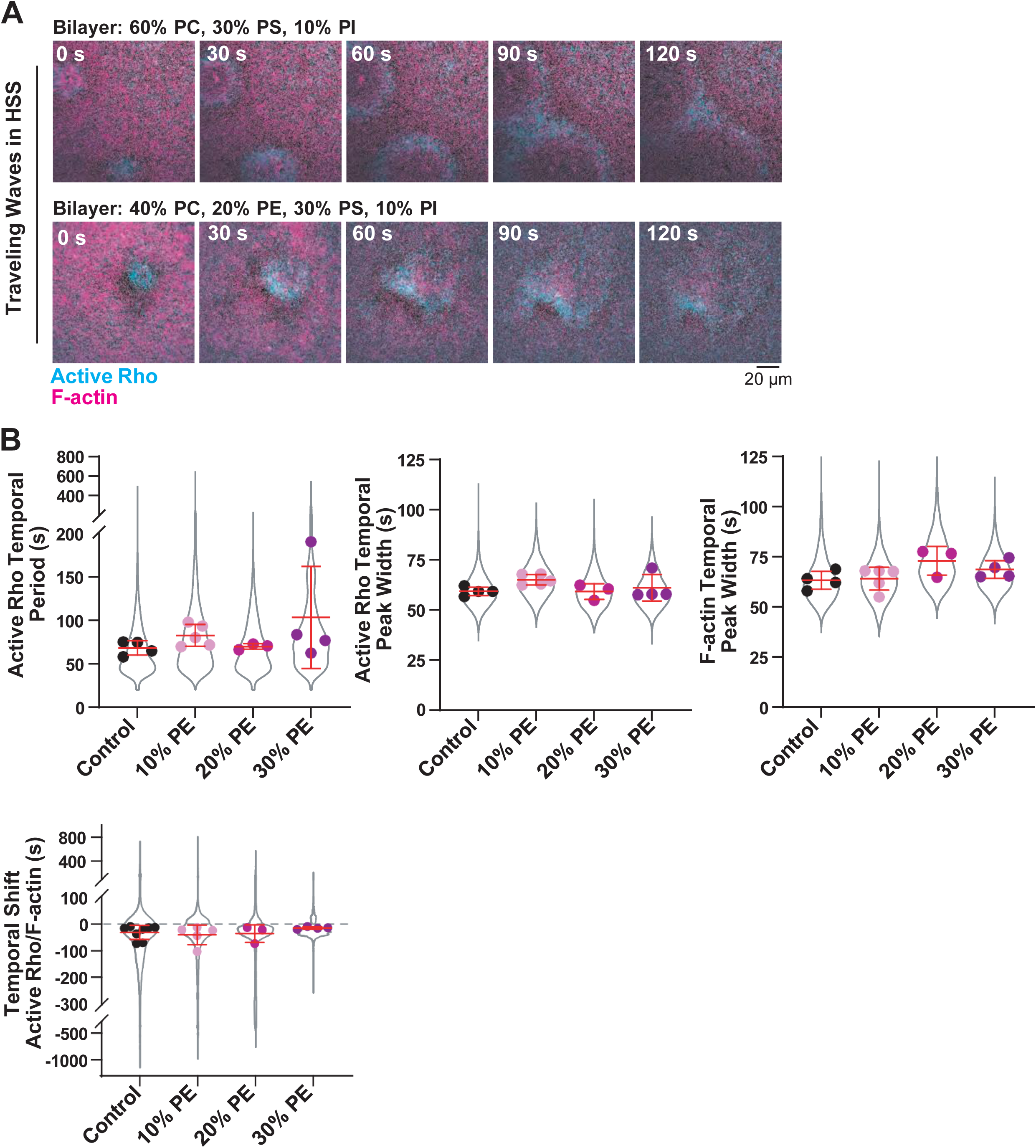
Supplemental analysis of active Rho and F-actin standing oscillations on PE-containing SLBs in HSS, related to Figure 4. A) Representative micrographs of traveling waves in HSS on SLBs with or without PE. Time is indicated in seconds after the initiation of traveling wave formation. B) Quantification of standing oscillations in HSS. Active Rho temporal period, active Rho and F-actin temporal peak width, and temporal shift on SLBs containing 0% (control, black), 10% (lavender), 20% (magenta), or 30% (purple) PE. Values from all experiments are represented by the violin plot (gray) and the mean of each experiment is shown as a dot. Red bars indicate total mean ± SD of the experimental means. For control, n = 3872 values from 4 videos from 3 biological samples from 3 experimental days. For 10% PE, n = 5099 values from 5 videos from 3 biological samples from 3 experimental days. For 20% PE, n = 3354 values from 3 videos from 3 biological samples from 2 experimental days. For 30% PE, n = 8222 values from 4 videos from 4 biological samples from 3 experimental days.

**Figure S4:**
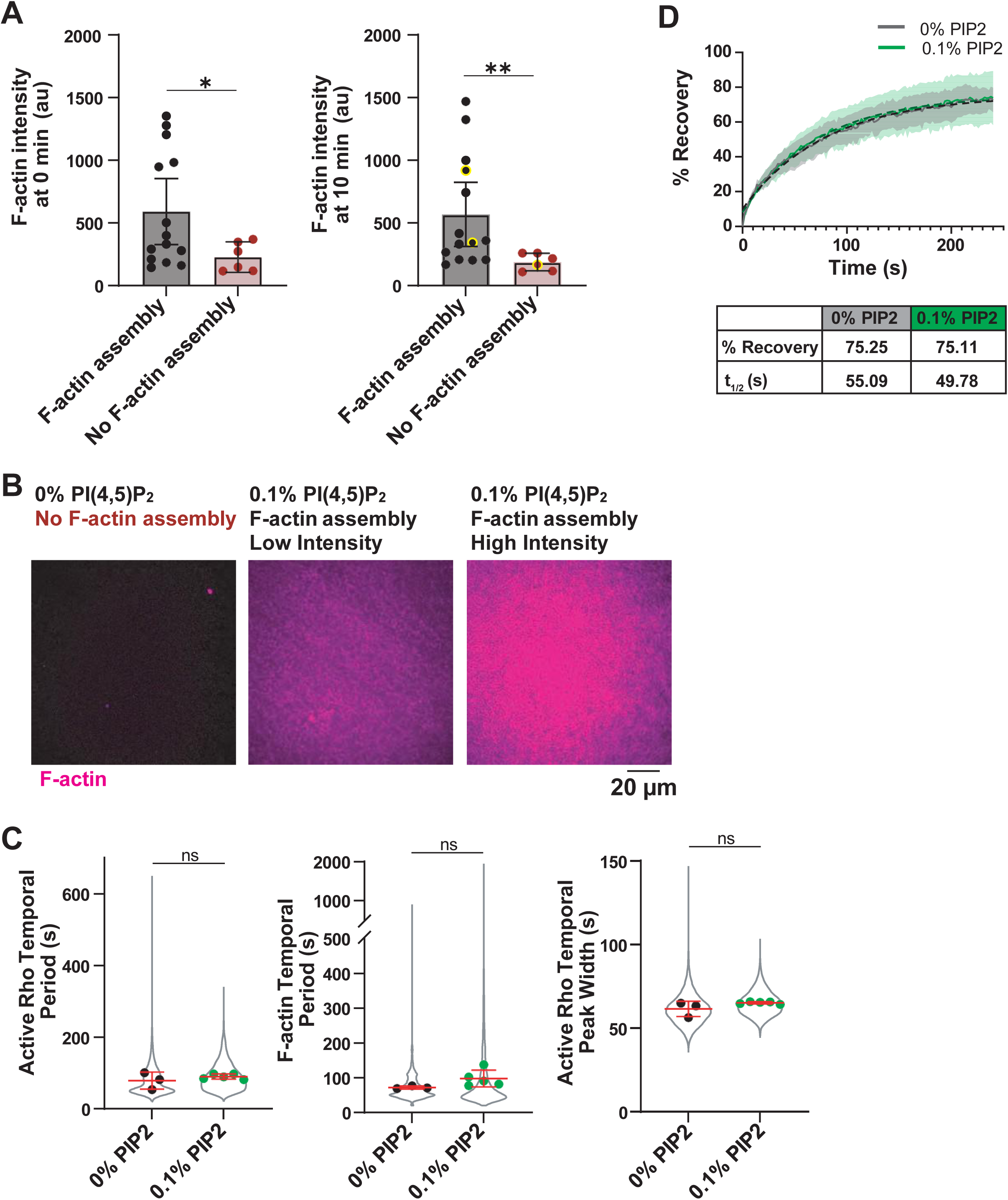
Supplemental analysis of F-actin intensity and cortical oscillation dynamics on 0 and 0.1% PIP_2_ SLBs, related to Figure 5. A) Analysis of F-actin whole field intensity on SLBs scored positive (gray) or negative (red) for F-actin assembly at the start of imaging (0 min, left) or 10 minutes after starting imaging (right). Mean ± 95% CI is shown. Yellow outline indicates data points that correspond to representative micrographs shown in (B). For F-actin assembly n = 14 videos, for no F-actin assembly n = 6 videos. * p ≤ 0.05, ** p ≤ 0.005. B) Representative micrographs of F-actin assembly on SLBs containing 0% or 0.1% PI(4,5)P_2_ 10 minutes after the start of imaging. Images correspond to data points highlighted in (A). Intensity is scaled equally across all images. C) Quantification of F-actin dynamics in HSS on SLBs containing 0% (black) or 0.1% (green) PIP_2_. Data points from all experiments are represented by a violin plot (gray) and the mean of each experiment is shown as a dot. Red bars indicate total mean ± SD of the experimental means. For 0% PIP_2_ SLBs, n = 3453 values from 3 videos from 3 biological samples from 2 experimental days. For 0.1% PIP_2_ SLBs, n = 5618 values from 5 videos from 3 biological samples on 2 experimental days. D) Fluorescence recovery after photobleaching (FRAP) of Cy5-labeled PC in SLBs containing 0% (gray) or 0.1% (green) PIP_2_. Mean (solid line) ± SD (shading) is shown. Dashed lines indicate the best fit one-phase association curve. For both, n = 7 bleach regions on 5 SLBs on 2 experimental days.

**Figure S5:**
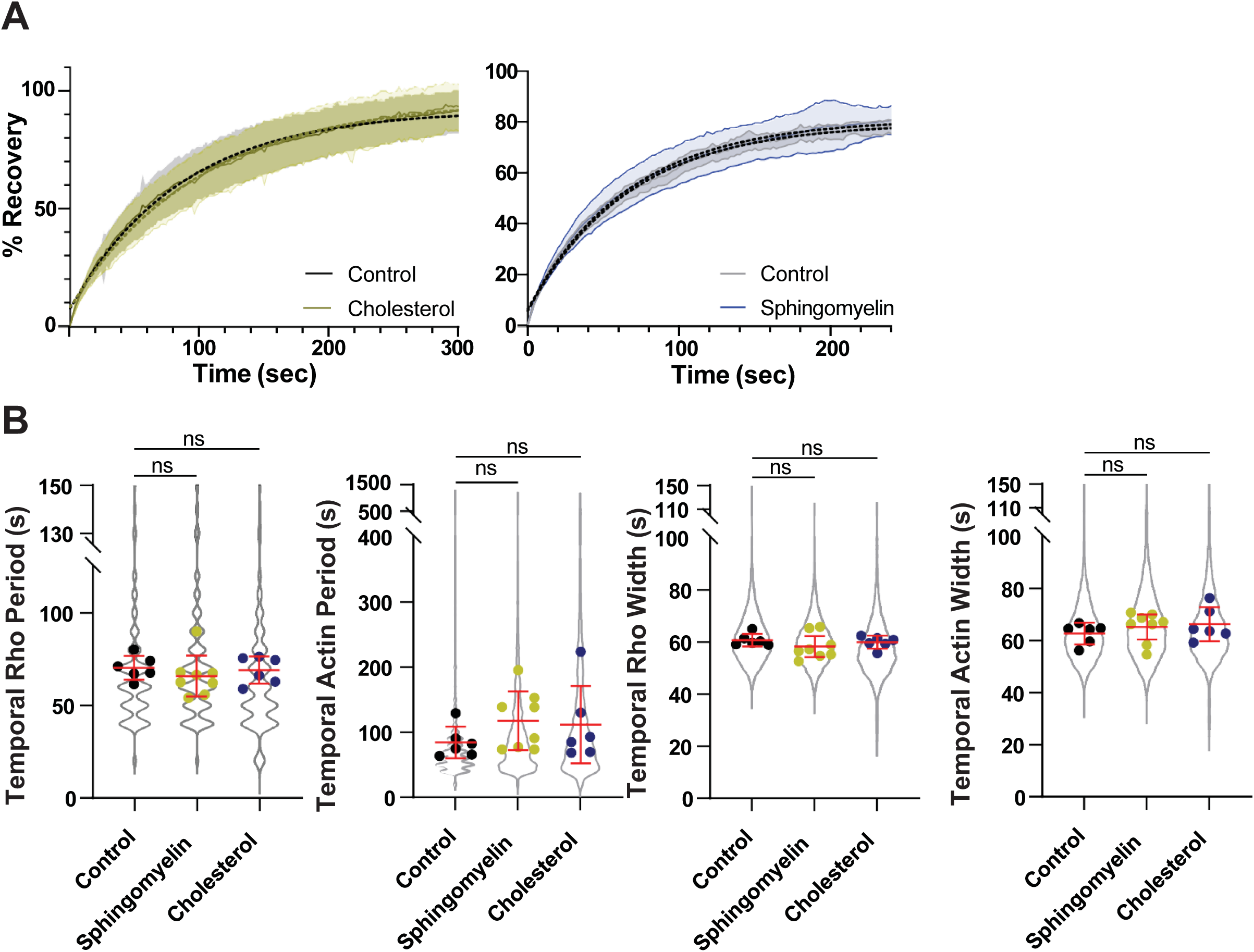
Supplemental analysis of active Rho and F-actin dynamics in LSS on cholesterol and sphingomyelin SLBs, Related to Figure 6. A) Intensity recovery after photobleaching curves of Cy5-PC labeled SLBs of the defined compositions (top). Control (gray), cholesterol-containing bilayers (yellow), and sphingomyelin-containing bilayers (blue) average recovery curves (solid line) ± SD (shading) is shown. Dotted line indicates the best fit one-phase association curves. For quantification of FRAP of cholesterol SLBs, n = 27 videos from 27 control SLBs across 6 experimental days and n = 18 videos from 21cholesterol SLBs across 4 experimental days were used. For quantification of FRAP of sphingomyelin SLBs, n = 6 videos from 3 control SLBs and 6 videos from 3 sphingomyelin SLBs across 2 experimental days. B) Quantification of the active Rho and F-actin temporal period and temporal peak width on control, sphingomyelin, and cholesterol-containing SLBs. All data points are represented in the violin plot and the average for each experiment is shown as a single dot. Red bars indicated the total average and standard deviation. For controls, n = 5623 values from 6 videos from 3 biological replicates across 4 experimental days. For sphingomyelin-containing SLBs, n = 6606 values from 6 videos from 3 biological replicates across 4 experimental days. For cholesterol-containing SLBs, n = 6647 values from 8 videos from 6 biological replicates across 3 experimental days.

## MATERIALS AND METHODS

### Experimental Model

Adult, wild type, *Xenopus laevis* females were injected with human chorionic gonadotropin (HCG) to induce egg laying. Frogs received a priming injection of HCG at least 2 days before inducing. Eggs were collected overnight in calcium-free MMR (Ca^2+^ free MMR: 2M NaCl, 40 mM KCl, 20 mM MgCl_2_, 2mM EGTA, 100mM NaHEPES, pH 7.8)

Frogs were fed twice per week with frog brittle (Nasco) and housed in a recirculating tank system (Aquaneering). The water quality parameters (temperature, pH, and conductivity) were constantly monitored for an optimal environment for frog health. Animal care staff monitored the frog health and water quality daily.

All studies strictly adhered to the compliance standards of the US Department of Health and Human Services Guide for the Care and Use of Laboratory Animals and were approved by Dartmouth College’s *<u>Animal Care and Use Committee</u>*. A board-certified Laboratory Animal Veterinarian oversees our animal facility.

### Experimental Methods

### Supported lipid bilayers

Powdered lipid stocks (Avanti Polar Lipids: Brain PC, 840053P; Brain PS, 840032P; Liver PI, 840042P; Brain PE, 840022P; 18:1 Cy5-PC, 850483C; Rhodamine-PE, 810150P; Brain PI(4,5)P2, 840046P; Brain Sphingomyelin, 860062P; Cholesterol, 700000P were resuspended in chloroform (Sigma-Aldrich, 650498) to a 20 mM stock concentration. Lipids were combined in the molar ratios of: 60% PC, 30% PS, and 10% PI (Stock A, Control); 60% PC, 30% PS, 5% PI, and 5% PI(4,5)P_2_ (Stock B); 50% PC, 10% Cy5 PC, 30% PS, 10% PI (Cy5-PC); 30% PC, 30% PS, 10% PI, 30% PE (PE), 30% PC, 30% PS, 10% PI, 20% PE, 1% Rhodamine PE (Rhodamine PE), 48% PC, 24% PS, 8% PI, and 20% cholesterol (cholesterol); or 45% PC, 25% PS, 7.5% PI, and 25% sphingomyelin (sphingomyelin). Lipids were dried down at 42°C with a steady stream of N_2_ gas on a heat block and then vacuum desiccated for 1 h at 45^°^C using a speed vac (Thermo Scientific, SPD1030). Lipid stocks were resuspended in high-salt *Xenopus* buffer (HS-XB: 4 M KCl, 20 mM MgCl_2_, 2 mM CaCl_2_) to 5 mM under N_2_ atmosphere, vortexed, incubated at 37^°^C for 30 minutes, and vortexed again. The lipid stocks were sonicated using a cup horn sonicator (QSonica) in polystyrene tubes at 80W for 5 mins with pulsed sonication (30s on; 10s off) to create small unilamellar vesicles (SUVs) under N_2_ atmosphere at room temperature. SUVs were filtered using 0.22 µm filter (Fisher Scientific, SCGP00525) into borosilicate glass vials (Thermo Scientific, 4010-17) under N_2_ atmosphere and stored in -20^°^C for less than 3 months.

SUVs were thawed at room temperature under N_2_ atmosphere to create supported lipid bilayers (SLBs). When needed, lipid stocks (listed above) were mixed to generate the desired ratio of PIP_2_ or fluorescently-labeled lipid. For example, in assays comparing active Rho and F-actin oscillations on cholesterol and sphingomyelin SLBs, PIP_2_ SUVs were combined with cholesterol or sphingomyelin SUVs to create 0.1% PIP_2_ SLBs. SUV solutions were sonicated under N_2_ atmosphere and diluted to 0.5 mM using HS-XB and CaCl_2_ was added to 4mM. CaCl_2_-treated SUVs were added to O_2_ plasma-cleaned imaging wells and incubated at 37^°^C for 30 minutes. The SLBs were washed three times with *Xenopus* buffer (XB: 100 mM KCl, 1mM MgCl_2_, 10 mM HEPES, pH 7.7) without exposing the SLBs to air. Washed SLBs were kept at room temperature for less than 4 hours before imaging experiments.

### Preparing Imaging Wells

24-well glass bottom chambers (Ibidi, 81817) were cleaned with 2% Hellmanex III (Sigma-Aldrich, Z805939) for 2 hours at 80^°^C with constant stirring. Wells were washed thoroughly deionized water and left to dry in a dust-free chamber. The wells were O_2_ plasma cleaned (Plasma Etch, 775-883-1336) for 10 minutes immediately prior to use.

Actin intact extract prep (CSF-arrested, M-phase, Low-speed supernatant [LSS]):

Adult, female *Xenopus laevis* were induced with HCG in Ca^2+^ free Marc’s Modified Ringers (Ca^2+^ -free MMR: 100 mM NaCl, 2mM KCl, 1mM MgCl_2_, 100 uM EGTA, 5mM HEPES, pH 7.8) to lay eggs. Eggs were de-jellied with 2% cysteine (Sigma-Aldrich, 168149) in Ca^2+^-free MMR at pH 7.8 for 10 minutes or until eggs jelly coat was removed. The eggs were washed twice with cytostatic factor *Xenopus* buffer (CSF-XB: XB + 1 uM MgCl_2_, 5 mM EGTA) while removing lysed and activated eggs (gardening). De-jellied eggs were washed in CSF-XB with a protease inhibitor cocktail (LPC: 10 µg/mL leupeptin, pepstatin A, and chymostatin; Sigma-Aldrich, L2884, 11524488001, C7268) and transferred to a thin-walled UltraClear centrifuge tube (Beckman Coulter) submerged in CSF-XB + LPC. The eggs were packed at 700 x g for 30 seconds followed by 1,400 x g for 15 seconds in a clinical centrifuge (Internation Equipment Co. Model CL). Excess buffer was removed after packing. The eggs were crushed in an ultracentrifuge (Beckman) using a swinging bucket rotor (SW55) at 15,000 x g for 15 minutes at 4^°^C. The cytoplasmic fraction was harvested using a needle and syringe (Fisher, 14955464; BD, 305196) through the side of the centrifuge tube while not disturbing the lipid or pellet fractions. Protease inhibitors (LPC) were added to a final concentration of 10 µg/mL with an energy mix with a final concentration of 9 mM (energy mix: creatine phosphate, 1mM ATP, 1mM MgCl_2_; Sigma, CRPHO, M8266). The extract is arrested in metaphase of meiosis II and kept on ice until use. This extract is referred to as low-speed supernatant (LSS).

### Converting to I-phase and contractility assay

To confirm high-quality M-phase arrest, 2 µL droplets of the LSS were added to 2 mL of mineral oil (Crystalgen, 24-107) in 35mm well dishes (Fisher Scientific, 3535) and monitored under a stereoscope (VWR, 89404-492). High-quality extract was visibly contracted in the mineral oil after 5 minutes. Non-contractile LSS was not used. CaCl_2_ (Fisher Scientific, AC349610000) was added to LSS to a final concentration of 0.4 mM, vigorously mixed, and incubated at room temperature for 5 minutes. LSS was put back on ice, to prevent further progression through the cell cycle, and assayed for contractility. Successfully cycled I-phase extract was not contractile after 5 min under oil.

### Preparation of actin Intact high-speed supernatant (HSS)

To create high-speed supernatant (HSS), the I-phase LSS was pipetted into a thin wall centrifuge tube (Beckman Coulter, 344090) and centrifuged with a swinging bucket rotor (SW55) in an ultracentrifuge for 90 minutes at 4^°^C at 90,000 x g. The cytoplasmic fraction was harvested from the centrifuge tubes using needle and syringe (Air-tite Products, MS12712), taking care to minimally disturb the lipid fraction. The extract was centrifuged again for 30 minutes with the same conditions. Afterwards, the cytoplasmic fraction was again harvested and stored on ice.

### Freezing LSS and HSS

LSS and HSS were gradually frozen in microfuge tubes in a cryofreezing chamber (Analytics Shop, NL51000001) in -80^°^C freezer overnight.

### Preparing reactions for imaging

Recombinant protein probes were added to LSS or HSS at the final concentrations: GFP-rGBD, 750 nM and UtrCH-594 or UtrCH-647, 100 nM. SLBs were washed once with extract, without exposing the SLB to air, to reduce dilution of cytoplasm on the bilayer for image acquisition. Extract with recombinant probes was added to washed SLBs and immediately transferred to the microscope for imaging.

### Drug Treatments

GSK-A1 (type III phosphatidylinositol 4-kinase PI4KA inhibitor, MedChemExpress) was resuspended in DMSO and stored at -80^°^C. HSS was treated with 60 µM GSK-A1 or an equivalent volume of DMSO as a vehicle control for 10 minutes at room temperature before adding to SLBs.

### Lipidomic analysis of LSS and HSS

Dalton Bioanalytics performed lipidomic mass spectrometry analysis of three biological replicates of LSS or HSS egg cytoplasm.

Sample Preparation and Lipid Extraction: LSS and HSS samples were stored at -80°C until lipid extraction. Sample processing order was randomized to minimize batch effects. Lipids were extracted by adding 80 μL of LC-MS grade isopropanol to 20 μL of sample, followed by vortexing and then incubation at 4°C for 30 minutes. Samples were then clarified by centrifugation at 14,000 × g for 5 minutes at 4°C, and the supernatant was transferred to LC-MS vials and stored at 4°C until analysis. Quality control (QC) samples were prepared by pooling small aliquots from each sample extract, with technical replicates generated from the pooled QC material. Water extraction blanks were included as negative controls.

### LC-MS/MS Analysis

#### Liquid Chromatography

Untargeted lipidomics analysis was performed using a Thermo Vanquish Flex system coupled to a Thermo Q Exactive Plus mass spectrometer. Chromatographic separation was achieved using a Waters Acquity CSH C18 column (1.0 × 150 mm, 1.7 μm particle size) maintained at 55°C. The injection volume was 5 μL with a constant flow rate of 0.070 mL/min over a 40-minute gradient. The mobile phase consisted of three components: (A) water with 0.1% formic acid, (B) acetonitrile with 0.1% formic acid, and (C) isopropanol with 5 mM ammonium acetate and 5 mM formic acid. The gradient program was as follows: 0-1 min, 96% A/2% B/2% C; 1-2 min, linear gradient to 0% A/98% B/2% C; 2-3 min, hold at 0% A/98% B/2% C; 3-15 min, linear gradient to 0% A/10% B/90% C; 15-28 min, hold at 0% A/10% B/90% C; 28-29 min, return to 0% A/98% B/2% C; 29-31 min, equilibration to 96% A/2% B/2% C; 31-40 min, hold at starting conditions.

#### Mass Spectrometry

Mass spectrometric analysis was performed using a Q Exactive Plus Orbitrap mass spectrometer operated in both positive and negative ionization modes with fast polarity switching. Data-dependent acquisition (DDA) was employed with Top7 fragmentation. Full MS parameters included: resolution of 70,000 FWHM, AGC target of 3×10⁶, maximum injection time of 100 ms, and scan range of m/z 175-1675. Data-dependent MS² parameters were: resolution of 17,500 FWHM, AGC target of 1×10⁶, maximum injection time of 50 ms, isolation window of 0.5 m/z, stepped normalized collision energy (NCE) of 20, 30, and 40, dynamic exclusion duration of 30 seconds, and intensity threshold of 1×10⁴. Isotopes were excluded from fragmentation, and charge states 2-8 and >8 were excluded from selection.

Data Processing and Lipid Identification: Raw LC-MS/MS data files were converted using MSConvert and processed using Dalton Bioanalytics’ custom R pipeline incorporating label-free quantification with match-between-runs (LFQ-MBR). Mass spectral data underwent retention time and m/z alignment with 5 ppm mass tolerance and 1-minute time tolerance. Lipid identification was achieved through two complementary approaches: (1) higher-confidence MS² spectral matching for tentative identifications using the Lipidex software, and (2) MS¹ accurate mass matching with predicted retention time validation based on the MS² putative identifications. Lipidomic profiles were normalized at the feature level using retention time-based loess recentering to a QC sample template. Lipid species were condensed to the isomeric species level (e.g., PC 35:1) by averaging across individual fatty acid sum compositions. All quantification was performed using relative abundance measurements.

#### Statistical Analysis

Differential lipid analysis was performed using limma linear modeling in R to compare low-speed versus high-speed centrifugation fractions. Each lipid feature was tested for association with the experimental grouping variable, with high-speed samples serving as the reference group. Quality control metrics included assessment of coefficient of variation (CV) in pooled QC samples, Pearson correlation analysis between QC technical replicates, and principal component analysis (PCA) to evaluate sample clustering. Lipid class enrichment analysis was conducted to identify systematically altered lipid categories using the fgsea R package. Statistical significance was assessed with false discovery rate (FDR) correction for multiple comparisons at FDR < 0.05. Data visualization included volcano plots, heatmaps, and retention time/mass-to-charge ratio distribution plots to assess differential abundance patterns across lipid classes. Note that the retention time-based loess normalization to QC samples may have dampened some of the observed differences between experimental groups.

### Live Imaging

For most of the experiments total internal reflection fluorescence (TIRF) microscopy was performed using a Nikon iLas Ring TIRF, with a Ti2 inverted, motorized microscope base, LUNF laser launch (488nm/561nm/640nm), perfect focus, and photo-stimulation module controlled by NIS Elements software. A TIRF Quad Dichroic cube (C-FL TIRF Ultra Hi S/N 405/488/561/638 Quad Cube, Z Quad HC Cleanup, HC TIRF Quad Dichroic, in metal cube, HC Quad barrier Filter) was used with a 60X Objective lens (CFI60 Apochromat TIRF 60X Oil Immersion Objective Lens N.A. 1.49, W.D. 0.12 mm, F.O.V 22 mm) and a Photometrics Prime 95B Back-illuminated sCMOS camera. For studies investigating cortical dynamics in LSS on PE-containing SLBs, imaging was performed using a Nikon Ti2-E motorized inverted microscope, without the ring TIRF module (point TIRF).

For investigating active Rho and F-actin dynamics, imaging started within 5 minutes of the addition of HSS with the recombinant protein probes to the SLBs. The SLBs were imaged in 10s intervals for 30-60 minutes in the appropriate channels with perfect focus on for the duration of the live imaging.

For the active Rho and F-actin polymerization assays in Figure 7, imaging wells were placed imaged on the TIRF microscope without extract (first 10 frames of videos). HSS with recombinant probes was added to the well while acquiring images and imaged for 10 minutes.

For FRAP experiments, 1.1 µm x 1.1 µm box with the ROI at the center of the field of view and bleached with a 405 nm laser for 1-2s at 15% laser power. The SLBs were imaged for 10s pre bleaching to get an average background intensity and for 5 minutes post bleaching.

### Quantification and Statistical Analysis

#### Image Pre-processing

Videos of active Rho and F-actin dynamics on SLBs were processed in Fiji (Schindelin *et al*., 2012) and cropped to the central portion of the field of view at approximately 152 µm^2^. Relevant frames were selected for analysis based on visible appearance of active Rho signal. Each channel was adjusted for visualization and pseudo-colored as indicated in figures. To remove non-dynamic signal, images were difference subtracted by 30 seconds using the “image subtraction” tool in Fiji or custom Fiji macro (diffMC, Zac Swider github (Swider *et al*., 2022)). Kymographs were created using the “Multi Kymograph” tool in Fiji from a line of width 5 pixels drawn across the video, as indicated on the figures or a custom Fiji macro (resliceMC, Zac Swider, github (Swider *et al*., 2022)). Kymographs were created from each channel individually and then merged.

### Quantification of Whole-field Intensity of active Rho and F-actin

Whole-field intensity was measured using the raw data (not difference subtracted) cropped to the central portion of the field of view (150 µm x 150 µm). Minimum and Mean gray values were measured for the respective channels over time using the “multi-measure” tool in Fiji. Mean values were normalized relative to the minimum intensity for each frame. Normalized mean intensity over time ± standard deviation was plotted in Prism (GraphPad).

### Scoring F-actin assembly on bilayers with and without PIP_2_

The assembly of F-actin was monitored for 10 minutes after extract addition to the bilayer. Bilayers were scored as “F-actin assembly” if linear F-actin arrays were clearly visible at the defined time points, regardless of the intensity. Bilayers with punctate or no F-actin signal were scored as “No F-actin assembly”. The quantification of F-actin intensity (Figure S2A) was performed as described for the “Quantification of Whole-field intensity” above, but for only two time points (0 and 10 minutes after addition).

### Traveling wave velocity analysis

Kymographs of active Rho and F-actin traveling waves were generated from difference subtracted movies using 1-pixel lines oriented parallel to the direction of traveling wave movement. Multiple traveling waves were measured for each movie. The F-actin channel was used to identify time points with continuous wave velocity and the rate was calculated as the change in distance divided by the change in time, using start and end points identified in the kymograph. For movies with repeated traveling waves only the first wave front was measured.

### Analysis of active Rho and F-actin dynamics

Difference subtracted videos were analyzed using waveanalysis (Landino et. al., 2021, Swider et. al., 2022) to quantify dynamics of active Rho and F-actin over time. The field of view was subdivided into boxes 4 µm x 4 µm in size. Boxes that produced no data (blank) for any one parameter (active Rho period, F-actin period, active Rho peak amplitude, F-actin peak amplitude, active Rho peak relative amplitude, F-actin peak relative amplitude, active Rho temporal width, F-actin temporal width, active Rho peak max, F-actin peak max, active Rho peak min, F-actin peak min, active Rho peak offset, F-actin peak offset, active Rho – F-actin shift, and active Rho-F-actin percent phase shift). were removed from the dataset. Videos with less than 50% total boxes removed were not used in our analysis. Data was also thresholded for a cross-correlation function that indicated active Rho signal leads F-actin (negative peak values).

### FRAP analysis

The ROI and background intensity was measured in Fiji by creating ROIs the size of the bleached region (1 µm x 1 µm); one in the bleached region and on outside the bleached region. The background intensity was used to correct for photobleaching Cy5-PC during image acquisition. The mean intensity of the ROI was calculated for all points starting at the first frame after photobleaching in time starting at the first frame imaged post photobleaching. Intensity measurements were background corrected using the following equation:

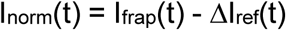

Where I frap is the ROI tracking the bleached area, I ref is the ROI tracking a region of the SLB outside the bleached area. Then values were normalized to a percent recovery using the following equation:

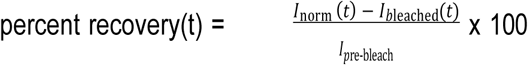

Where I_bleach_ is the intensity immediately after bleaching and I pre bleach is the average intensity of the ROI before bleaching. The average % recovery ± SEM was plotted using GraphPad Prism. The mean percent recovery of the as fit to a single exponential association curve using the ‘‘analyze’’ function in GraphPad Prism. The curve was constrained so that the plateau was ≤ 100 and the Y0 was set to equal 0. All data points outside of this range were excluded from analysis. The SEM was calculated using equal variance.

Percent area of active Rho:

The mean area of the active Rho in a difference subtracted movie was analyzed for each frame of the movie by creating a mask using an intensity threshold of 50 arbitrary units. Total percent area and cluster size was calculated in Fiji.

### Active Rho binding and F-actin Polymerization Assay

For the active Rho and F-actin polymerization assays in Figure 7, the intensities were normalized by dividing the intensity of the current frame by the average of the last 10 frames of the movie using the following equation:

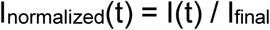

Where I_normalized_(t) is the normalized intensity, I(t) is the current intensity, and I_final_ is the average intensity of the final 10 frames of the video.

### Expression and Purification of recombinant protein

For recombinant protein expression, pFastBac1 constructs encoding FLAG-Cys-UtrCH and FLAG-GFP-rGBD were used to generate recombinant bacmids in DH10BAC (Invitrogen) bacteria. Sf9 infected cells were transfected with recombinant bacmids with PEI transfection reagent (Polysciences, 3966) in collaboration with bioMT core facility at Dartmouth. The virus was amplified to improve titter, and Sf9 cells were infected for three days at 27^°^C for protein expression.

Cell pellets were lysed in PBS + 0.1% TX-100 with protease inhibitors (LPC, PMSF, Sigma). Recombinant proteins were purified using anti-FLAG M2 affinity resin (Sigma) and arginine-based elution (Bement *et al*., 2015). Elution fractions were pooled and the protein was concentrated using Amicon Ultra Centrifugal Filters, 30 kDa MWCO (Millipore) and stored in 25 mM HEPES, 100 mM KCl, pH 7.5. FLAG-Cys-UtrCH was labeled using Alexa Fluor 594 Protein Labeling Kit or Alexa Fluor 647 Protein Labeling Kit according to the manufacturer’s instructions (ThermoFisher).

